# ERM-Dependent Assembly of T-Cell Receptor Signaling and Co-stimulatory Molecules on Microvilli Prior to Activation

**DOI:** 10.1101/766196

**Authors:** Shirsendu Ghosh, Vincenzo Di Bartolo, Liron Tubul, Eyal Shimoni, Elena Kartvelishvily, Tali Dadosh, Sara W. Feigelson, Ronen Alon, Andres Alcover, Gilad Haran

## Abstract

T-cell surfaces are covered with microvilli, actin-rich and flexible protrusions. We use super-resolution microscopy to show that ≥90% T-cell receptor (TCR) complex molecules TCRαβ and TCRζ, as well as the co-receptor CD4 and the co-stimulatory molecule CD2 reside on microvilli of human T cells. Furthermore, TCR proximal signaling molecules involved in the initial stages of the immune response, such as the protein tyrosine kinase Lck and the key adaptor molecule LAT, are also enriched on microvilli. Notably, phosphorylated proteins of the ERM (ezrin, radixin, moesin) family colocalize with these heterodimers as well as with actin filaments within the microvilli of resting T cells. This finding implies a role for one or more phosphorylated ERMs in linking the TCR complex to the actin cytoskeleton within microvilli. Indeed, expression of a dominant-negative ezrin fragment effectively redistributes TCR molecules over the whole T cell surface. Our results establish microvilli as key signaling hubs, in which the TCR complex and its proximal signaling molecules and adaptors are pre-assembled prior to activation in an ERM-dependent manner. The preformed positioning of these actin-binding TCR assemblies on individual microvilli can facilitate the local transmission of TCR signals seconds after TCR occupancy and impacts the slower subsequent events that lead to the assembly of immunological synapses.

## Introduction

T cells play a pivotal role in adaptive immunity. When encountering antigen-presenting cells (APCs), these cells undergo a major reorganization of their surface, with a subset of these encounters leading to the formation of immunological synapses (IS) (1–3). Recent evidence suggests that within seconds of T-cell receptor (TCR) occupancy by cognate antigenic peptides, the TCR complex transmits T cell activation signals in the form of phosphorylation and calcium mobilization (4–6). These signals are often generated by T cells that engage their APCs for minutes and even shorter times (4–6). These observations suggest that most TCR signaling events occur during brief contacts between the T cell and various APCs. How such short-lived contacts transmit T cell activation signals so efficiently remains an enigma. The TCR complex (composed of the transmembrane proteins TCRαβ, CD3γ,δ,ε and TCRζ) initiates the T-cell immune response after it recognizes foreign antigen-derived peptides bound to major histocompatibility complex (MHC) molecules on the surface of APCs. The roles of the TCR co-receptors, CD4 or CD8, adhesion and co-signaling receptors like CD2, and proximal signaling proteins such as the protein tyrosine kinase Lck, the adaptor LAT and the tyrosine phosphatase CD45 are well described (1, 2, 7, 8). However, how these molecules participate in translating the very initial TCR ligand occupancy events to productive TCR signaling is still poorly understood. Furthermore, while the involvement of multiple cytoskeletal elements in the formation of the IS has been documented (9–13), the role of cytoskeletal components in the initial antigen recognition events triggering TCR signaling remains unclear. While several T-cell related or more general models were proposed to explain protein distribution on cellular surfaces, (14–18) much remains to be learned about the organization of key surface receptors implicated in T-cell activation and the formation of functional immune synapses.

Recent advances in imaging techniques, such as fluorescence super-resolution microscopy, have led to the observation that different membrane receptors indeed exist as nanoclusters of an average size of a few tens to hundreds of nanometers even in non-activated T-cells. For example, Davis and co-workers suggested that in resting murine T cells, two very important TCR signaling proteins, TCRζ and LAT, pre-cluster into separate protein islands on the T-cell membrane (17). They also showed that in murine T cells, the co-receptor CD4 forms distinct nano-clusters (18). Using confocal, total internal reflection fluorescence, and superresolution microscopy, Rittera et al. unraveled the role of dynamic regulation of the cortical actin cytoskeleton in initiation and termination of secretion of “lytic granules”.(19) Samelson and coworkers reported that the kinase ZAP-70 co-localizes with TCRζ, while both partially mix with LAT molecules in human lymphocytes and Jurkat T cells (20). Gaus and coworkers showed that the active conformation of the key TCR associated kinase Lck promotes its clustering on the membrane of Jurkat cells, whereas the inactive conformation of this kinase hinders its clustering (21). They also showed that T-cell signaling initiation occurs only within dense clusters of TCR-CD3 complexes in Jurkat T cells (22). Finally, Lillemeier and coworkers demonstrated nanometer-scale clustering of TCR molecules on T cells in resting lymph nodes of mice (23).

Strikingly, all of these studies considered the T-cell membrane as a two-dimensional sheet, and ignored the complex 3D microvillar structures on the surface of T cells (24–26). These finger-like protrusions, ∼100 nm in radius and a few hundreds of nanometers in length, are highly flexible and dynamic. We recently discovered that in human peripheral blood T cells and in effector T lymphocytes differentiated from these lymphocytes, TCRαβ molecules are preferentially localized to microvilli, suggesting a role for these dynamic protrusions in initial T cell recognition of cognate antigenic peptides presented by dendritic cells and other APCs (11). Indeed, Krummel and co-workers demonstrated that microvilli function as sensors that scan the surface of encountered APCs within seconds (27). Sherman and co-workers also observed that microvilli are involved in early T-cell activation (12), and Jun and coworkers demonstrated that upon TCR activation T cells release nanoscale microvilli-related vesicles that interact with APCs and transfer cargo to these cells through a process termed trogocytosis (28).

Although these recent studies invoked a key role for microvilli in the ability of T cells to encounter antigenic peptides, they did not resolve how TCR-associated signaling molecules, which must operate shortly after the initial occupancy with a ligand, get rapidly recruited by stimulated TCR complexes. To shed light on this issue, we study here the distribution of key TCR signaling components both on human T effector cells and on Jurkat T cells, a prototype CD4 T cell line. (29) We find that all of these molecules (apart from CD45) are pre-segregated on the microvilli of both types of T cells. We further determine the mechanism of this unique enrichment of TCR signaling machineries on microvilli by interfering with their anchorage to the dense cortical actin cytoskeleton of the microvilli. Our work establishes microvilli as important hubs for TCR signaling, on which all key molecular players required for the initial T cell-APC interaction are pre-organized.

## Results

### Mapping the distribution of membrane proteins in relation to microvilli

We previously demonstrated that a combination of variable-angle total internal reflection microscopy (VA-TIRFM) and stochastic localization nanoscopy (SLN) can map the distribution of peripheral blood human T-cell surface proteins with respect to microvilli (11). Here we adopted the same methodology to study a large set of membrane proteins on the surface of resting, human effector T cells (Fig. 1A) as well as non-stimulated Jurkat cells (Fig. 1B). Each membrane protein of interest on the cells was labelled with an Alexa-647 tagged antibody. The cells were then fixed and stained with the dye FM143FX. A home-built TIRF microscope was used to image the cells, which were placed on a poly-L-lysine (PLL) coated surface (Figs. 1C-E). Since the cells were already fixed, the interaction with the PLL surface did not cause any distortion of their topography (See Supporting Text and Fig. S1), as directly verified by SEM (Figure 1A-B). Initially, VA-TIRFM images were obtained from treated cells residing on a glass surface. The fluorescence intensity measured at each point in an image was converted into the distance of that point from the glass (δz) (30, 31). The set of δz values obtained from VA-TIRFM images permitted us to plot a map of the membrane structure, thus obtaining the 3D membrane topography (Figs. 1F-G). By combining measurements at several angles of incidence of the excitation beam, we improved the statistical precision of the procedure (Fig. S1A-H).

**Fig. 1:**
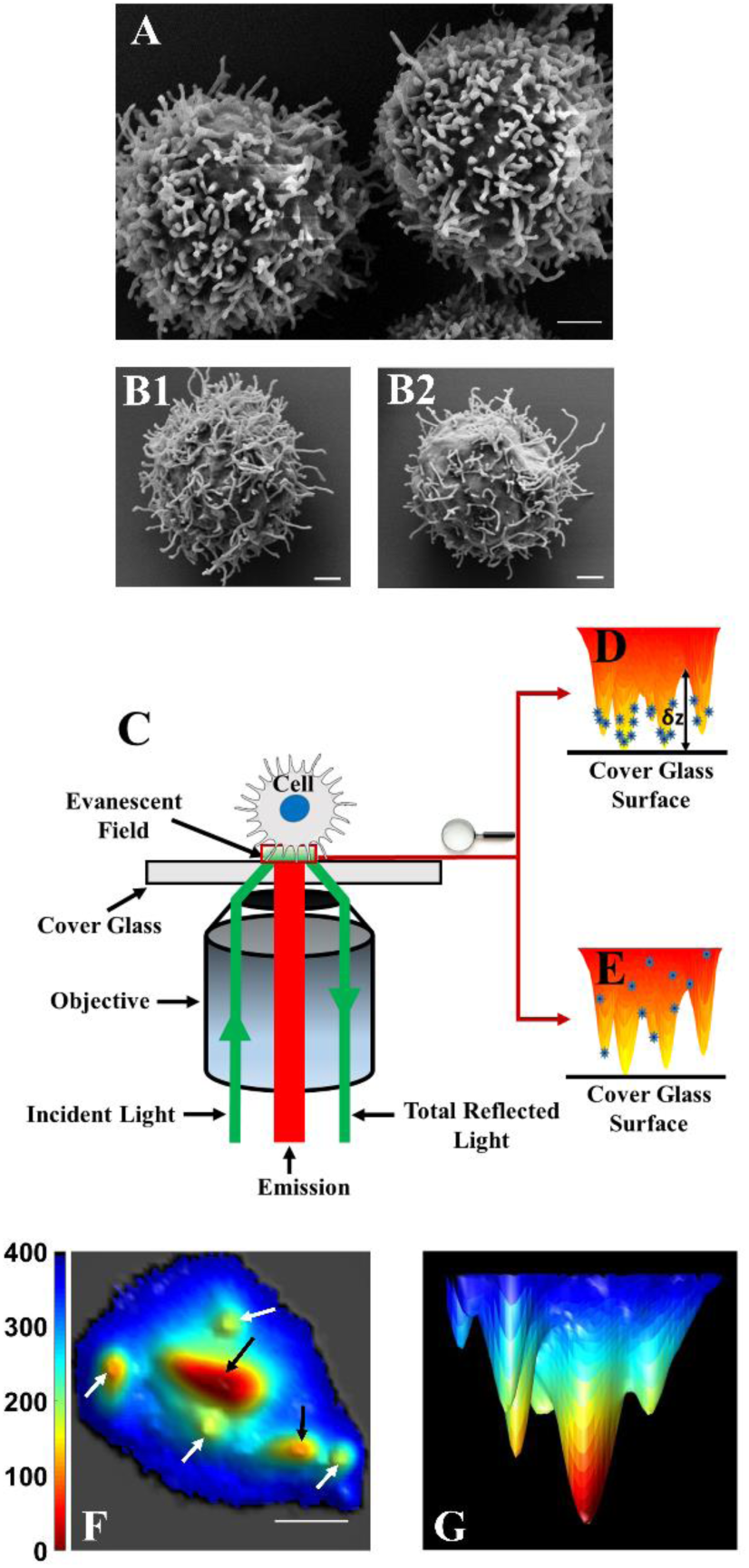
Combining VA-TIRFM and SLN for localizing molecules with respect to microvilli. (A-B) SEM images of T cells at a magnification of 20,000×, demonstrating the dominance of microvilli on the cell surface. (A) Human effector T cell. (B) Two examples of Jurkat T cells. Scale bars: 1 μm. (C) A schematic configuration of the TIRFM setup. (D) Schematic of a cell with proteins on the microvilli. The distance from the glass to the surface of the cell at each point (δz) is obtained from analysis of TIRF images measured over a range of angles of incidence. (E) Schematic of a cell with proteins randomly distributed throughout the membrane. (F) Bottom view of a representative 3D reconstruction map of a Jurkat cell membrane (white arrows indicate microvilli positioned vertically with respect to the glass surface and black arrows indicate microvilli lying down horizontally. Scale bars: 1 μm. (G) Side view of the 3D reconstruction map of the same Jurkat cell membrane shown in F.

The typical width of microvilli, as observed in SEM images (Fig. 1A-B), is 100 nm or less. This is clearly below the optical diffraction limit, and hence microvilli would appear thicker than in reality in any imaging technique that is not able to break this limit. TIRFM can obtain sub-diffraction resolution only in the Z direction, but is diffraction-limited in the X-Y plane, and this is why the microvilli appear in our TIRF images thicker than in SEM (Fig. S1I). Nevertheless, the density of microvilli calculated from segmentation maps of VA-TIRFM images compared well with the density obtained from SEM images (Fig. S1A-H), indicating that the method did not miss a significant number of microvilli due to the lower X-Y resolution. As an additional means for the assignment of observed structures as microvilli (especially elongated structures classified as microvilli lying horizontally), we performed L-selectin localization experiments (See Supporting Text and Fig. S1J-L).

We used SLN to obtain maps of antibody-labelled membrane proteins of interest. In SLN, the location of single emitters is determined with accuracy well below the diffraction limit (32–34). By superimposing the super-resolved membrane protein map on the 3D membrane topographical map, we were able to characterize the distribution of each protein molecule in relation to microvilli. We collected super-resolution images from two separate planes of each cell - right at the interface with the glass substrate (where mostly microvillar regions and a small fraction of cell body regions were observed, 0 nm plane) and 400 nm away from the substrate (where both microvilli and cell body regions were imaged, −400 nm plane). We verified that molecules could be detected with equal probability from the two planes by carrying out experiments on samples in which dye molecules were randomly distributed on either plane (see Supporting Text and Fig. S1M-O).

Using the two sets of images and detailed image analysis, we segmented the membrane area of each cell into either the microvillar region or the cell-body region, and then determined the distribution of each of the studied molecules in the two regions. We also quantified changes in the molecular distribution as a function of distance from the tips of microvilli (which were defined as those regions that are not more than 20 nm away from the pixel with the minimum δz value). More details about our image analysis methodology are provided in the SM, Materials and Methods section. However, it is important to point out here that this analysis does not depend on the diffraction-limited resolution of the TIRF images in the X-Y plane, but only on the high resolution in the Z direction, which is due to the properties of the evanescent field in TIRFM. We applied this methodology to study a series of TCR associated proteins on human effector T cells and Jurkat cells (Fig. 2 and 3). These experiments are described in detail below. Statistics of the number of cells measured in each experiment, as well as the percentage of molecules of different signaling protein localized in the microvilli, are provided in Supporting Materials (SM), Table S1.

**Fig. 2:**
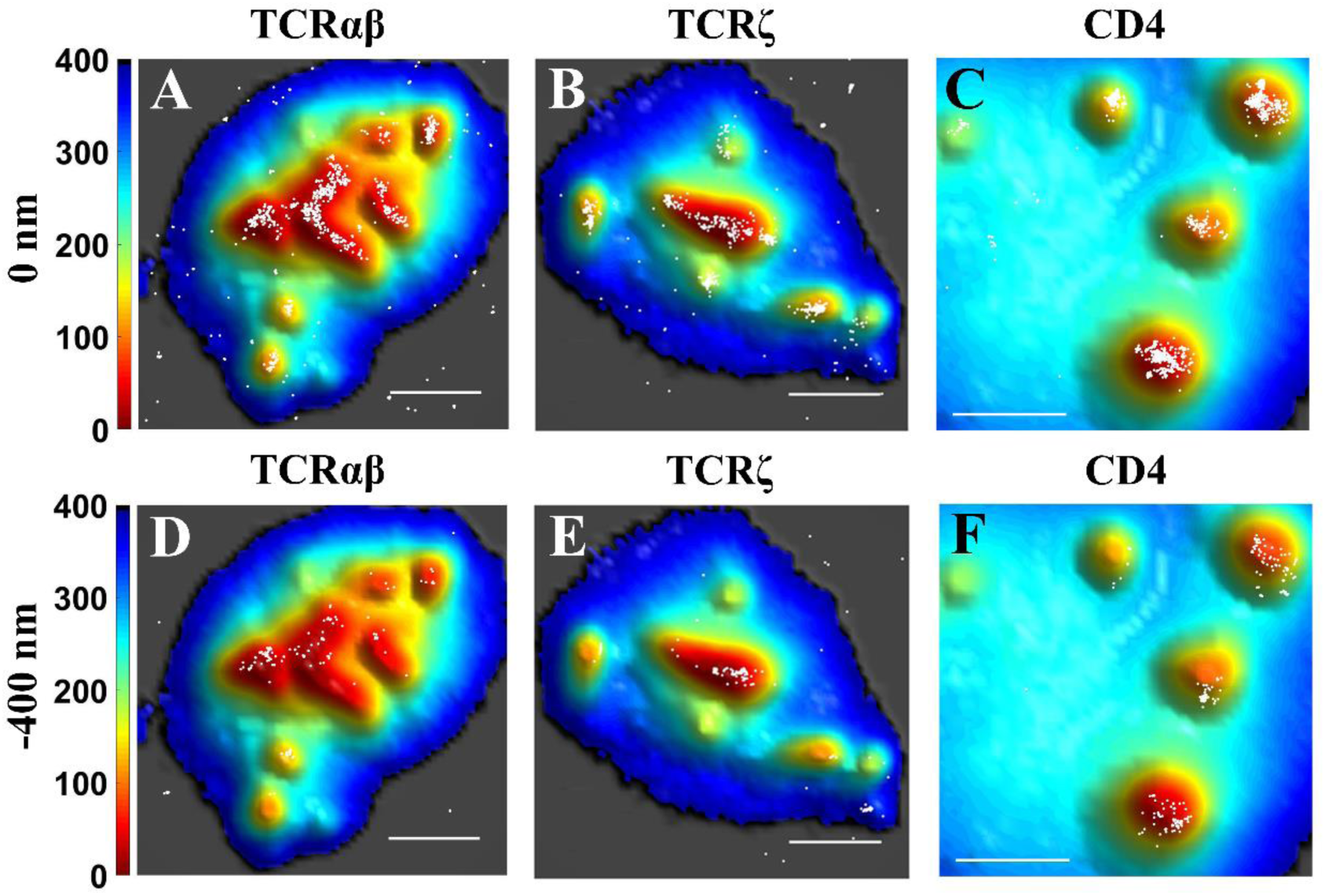
T-cell receptor related proteins are localized to the microvilli. Positions of protein molecules obtained from SLN (white dots) are superimposed on membrane topography maps of Jurkat T cells obtained from VA-TIRFM. The labels ‘0 nm’ and ‘-400 nm’ in panel titles refer to the two focal planes sampled in the super-resolution experiments. Scale bars: 1 μm.

**Fig. 3:**
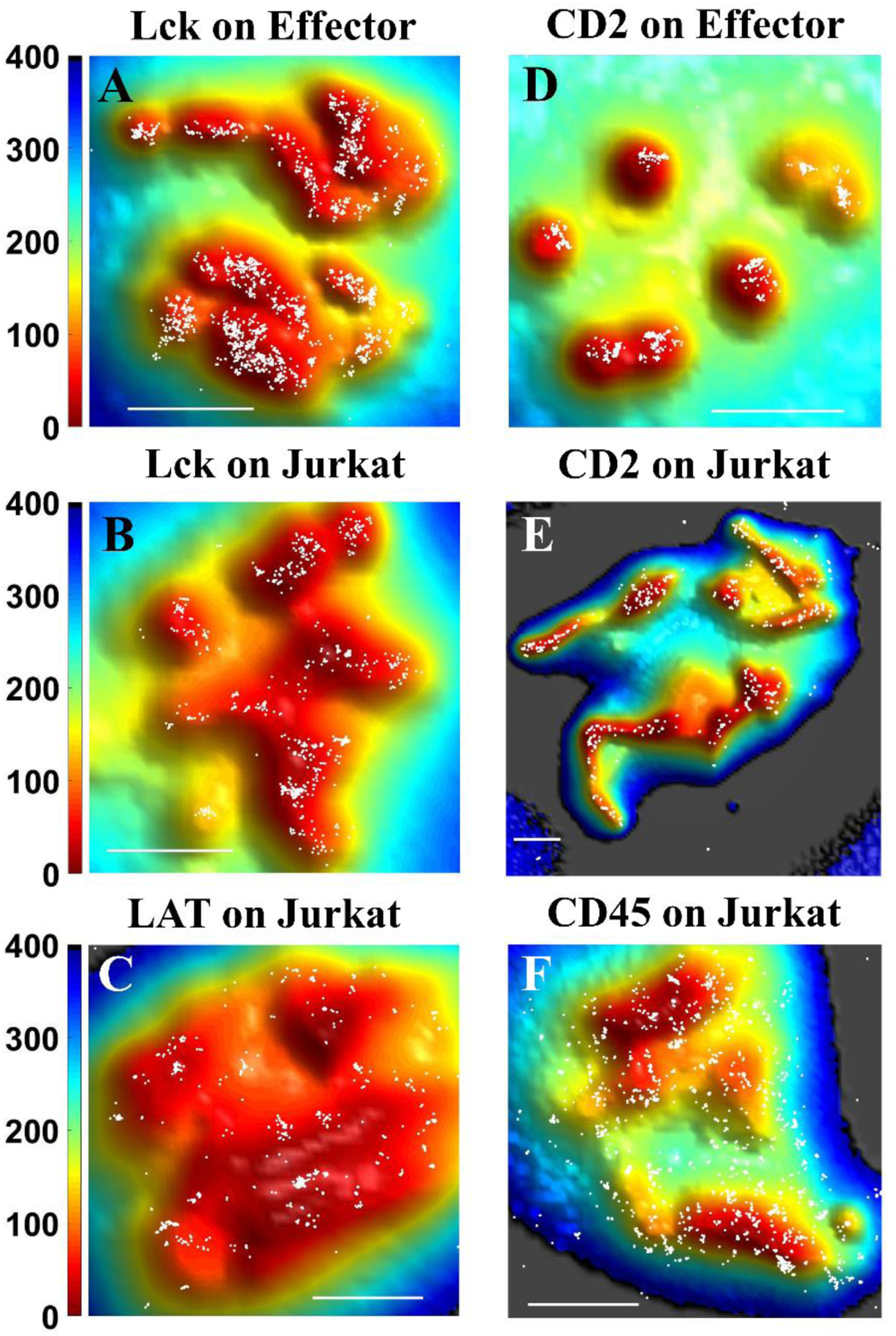
Localization maps of membrane proteins involved in the initial T-cell immune response. Positions of protein molecules obtained from SLN (white dots) at the 0 nm plane on either Jurkat T cells or human effector T cells are superimposed on membrane topography maps obtained from VA-TIRFM. The color bars represent distance from the glass in nm. Scale bars: 1 μm.

#### TCRαβ

TCRαβ is a dimer that constitutes the recognition unit of the TCR complex (35). It combines with CD3 and TCR**ζ** to carry out signal transduction upon binding to peptide-antigen-MHC molecules on the APC (35). In our previous work we showed that TCRαβ molecules localize almost exclusively to the microvillar regions of human peripheral T lymphocytes (11). We now verified this finding in Jurkat T cells. Indeed, we observed that TCRαβ molecules strongly localize to the microvillar regions of these cells as well (Fig. 2A,D). Our calculations showed that 89.0±2.4% of TCRαβ molecules reside on microvilli (Fig 4A). As shown in Fig. 4B, as the distance increases from the microvillar tip regions, a plot of the surface-normalized increase in the fraction of total TCRαβ molecules (δCount/δArea plot) falls sharply, pointing to strong localization on the microvilli. Cluster analysis using Ripley’s K function showed that the maximum cluster size of TCRαβ molecules is as high as 167±21 nm (Fig 4C, for a brief description of the analysis see Methods section), matching the typical radius of a microvillus.

**Fig. 4.**
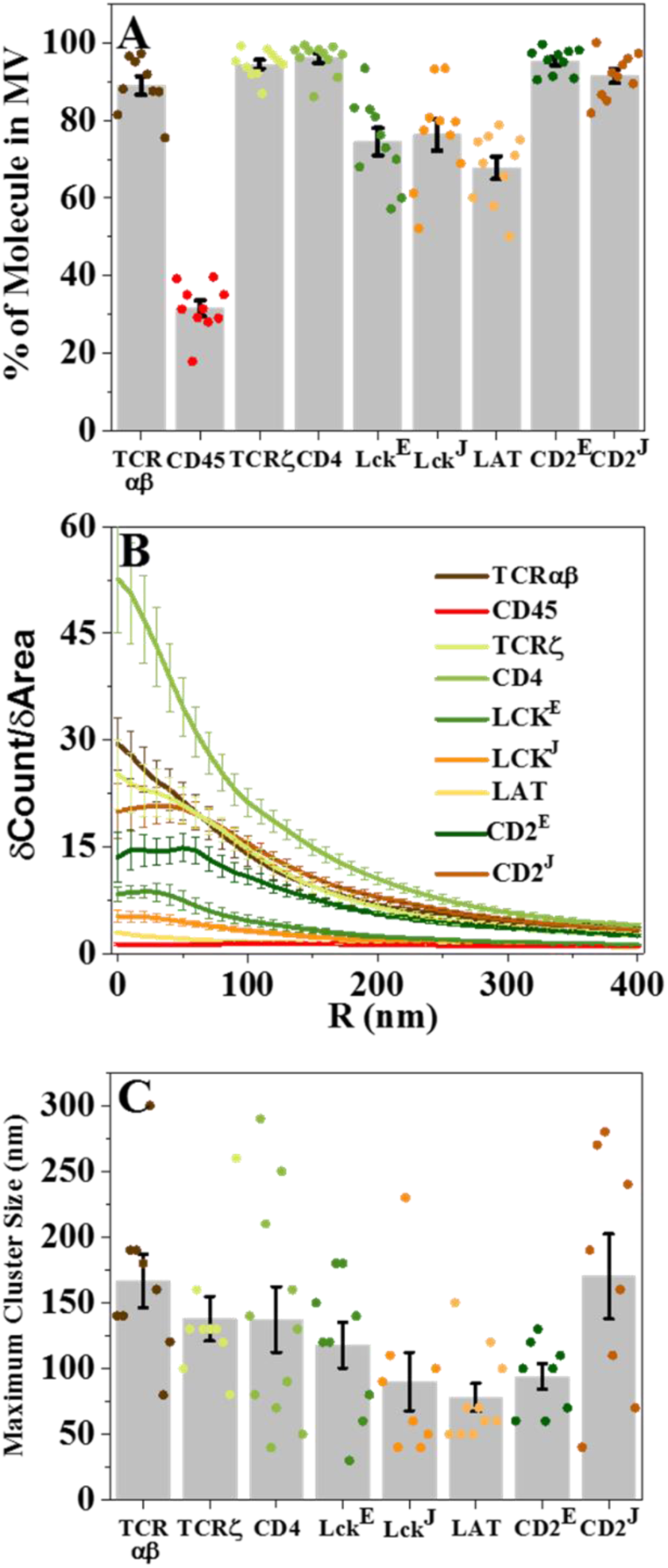
Quantitative measures for the distribution of proteins on the T-cell surface: (A) Percentage of molecules on microvillar (MV) regions of the membrane. The values for individual cells are shown as dots in the plot. (B) Cumulative increase of the fraction of total molecules on each cell as a function of the distance from the central microvilli region, normalized by the cumulative increase in the fraction of area (δCount/δArea) as a function of distance from microvilli. (C) Maximum cluster sizes. The values for individual cells are shown as dots in the plot. The gray bars are averages over all cells measured at the 0 nm plane. Error bars represent standard errors of the mean. The E and J superscripts following some protein names denote values obtained with effector cells and Jurkat cells, respectively. When no superscript is shown, the values were obtained with Jurkat cells.

#### TCRζ

TCRζ (also known as CD247 or CD3ζ) appears as a dimer, and is essential for the formation of the TCR complex and its stability (35). During the T-cell activation process, it is phosphorylated by Lck on its ITAMs (immune receptor tyrosine-based activation motifs), which facilitates interaction with ZAP70 (35). We observed that, like TCRαβ, TCRζ molecules reside almost exclusively on the microvilli of Jurkat T cells (Fig. 2B, E). Indeed, 96.0±1.2% of the molecules were localized to microvilli (Fig. 4A and SM, Table S1), and the δCount/δArea plot was very steep (Fig. 4B). The maximum cluster size of TCRζ receptors was 138±17 nm (Fig. 4C), again matching the microvillar radius.

#### CD4 (Cluster of differentiation 4)

This TCR co-receptor is a glycoprotein universally found on the surface of helper T cells and subsets of dendritic cells (18, 36, 37). It assists the TCR complex in forming connection to the antigen-presenting cell by binding MHC class II molecules, and it also recruits the key Src kinase Lck to the TCR. Importantly, Jurkat T cells display only CD4 and not CD8 on their surface (38).We found that 94.4±1.1% of the CD4 molecules are localized to the microvilli of Jurkat T-cells (Fig. 2C, F, see also Fig. 4A, SM, Fig. S2 and SM, Table S1). The slope of the δCount/δArea plot (Fig. 4B) was the largest among all the proteins measured. The maximum cluster size calculated for CD4 was 137±25 nm (Fig. 4C).

#### Lck (lymphocyte-specific protein tyrosine kinase)

Following engagement of the TCR by an antigen on the APC, Lck phosphorylates the intracellular domains of CD3 and TCRζ and later on Lck and TCR co-localize in the center of the IS (39). Lck was also found in CD2 clusters of activated T-cells (39). We observed that in both human effector T cells (Fig. 3A) and Jurkat cells (Fig. 3B), Lck molecules localize to a large extent to microvilli (for localization at the −400 nm plane see SM, Fig. S2A and B). 74.5±3.5% of all Lck molecules were localized on the microvillar regions of human effector T cells based on the 0 nm plane images, whereas 65.4±4.7% of the molecules were localized on the microvillar regions based on the −400 nm plane images (Fig. 4A and SM, Fig. S2G). A similar picture arose in the case of Jurkat cells, with 76.3±4.1% of all Lck molecules localized on microvilli regions based on 0 nm plane images, and 59.6±4.8% of the molecules localized on microvilli regions based on the −400 nm plane images (Fig. 4A and SM, Fig. S2G). These fractions are, however, significantly smaller than those of TCRαβ, TCRζ and CD4. Thus, although interacting with CD4 molecules (40), it seems that not all Lck molecules are directly linked to these co-receptor molecules, as also suggested by others studies (41).

#### LAT (Linker for activation of T cells)

This adaptor protein is phosphorylated by ZAP70 following activation of T cell receptors, and initiates a cascade of events involving multiple signaling molecules that leads to the formation of the IS.(35) Like Lck, LAT resides in intracellular vesicles exchanging with the plasma membrane.(35) We found that although LAT molecules were distributed throughout the Jurkat T-cell membrane, they still showed a significant preference for microvilli (Figs 3C and 4A, for the localization map and analysis of LAT at the −400 nm plane see SM, Fig. S2C and S2G-H). The fraction of LAT molecules localized on microvilli at the 0 nm and −400 nm plane images was 67.8±2.9% and 54.6±2.3%, respectively (Fig. 4A and S2G). The initial slope of the δCount/δArea plot (Fig. 4B) was not as high as for other membrane proteins, but still significantly larger than that of CD45 (SM, Table S2). Interestingly, LAT clusters tended to be smaller than those of TCR components (maximum size 78±10 nm, Fig. 4C), but their size agreed well with previous studies. (17, 42)

#### CD2 (Cluster of differentiation 2)

CD2 is an adhesion molecule that binds LFA-3 (CD58) on the APC membrane, and recruits essential kinases such as phosphoinositide 3-kinase to the vicinity of the TCR signaling complexes during the initial stages of the immune response, thereby amplifying TCR signaling (8, 43, 44). We found that CD2 molecules are also highly enriched on microvilli of human effector T cells (Fig. 3D and S2D) and Jurkat cells (Fig. 3E and S2E), with more than 90% of the molecules residing on these projections (Fig. 4A; For localization at the - 400 nm plane see SM, Fig. S2G.) The δCount/δArea plot is flat for in the first ∼50 nm but then decays (Fig. 4B), also pointing to the essentially exclusive residence of these molecules on microvilli. The maximal cluster size of CD2 molecules on the microvilli of effector and Jurkat cells was 94±10 and 170±32 nm, respectively (Fig. 4C). Thus, following TCR occupancy by ligand, LFA-3 occupancy of CD2 molecules may take place in the vicinity of the TCR, and instantaneously amplify the TCR signal.

#### CD45

CD45 is an abundant protein tyrosine phosphatase. This transmembrane protein has been shown to operate as a modulator of the sensitivity of T cells to activation, by associating with several membrane proteins, including the TCR complex, CD4/CD8, Lck and ZAP-70 (14, 35). In accordance with our previous finding in human peripheral T lymphocytes (11), we found that CD45 also distributed throughout the membrane of Jurkat T cells (Fig. 3F, for localization at the - 400 nm plane see SM, Fig. S2F). Only 31.6±2.0% of CD45 molecules localized in microvilli regions (Fig. 4A). The near-zero slope of the δCount/δArea plot for CD45 (Fig. 4B) also pointed to a homogeneous distribution of the CD45 molecules all over the membrane. Interestingly, two recent studies also demonstrated a uniform distribution of CD45 on the T-cell membrane (27, 45).

### The actin cytoskeleton and localization of membrane proteins on microvilli

The role of the actin cytoskeleton in receptor clustering and in IS formation has been extensively studied (3, 19, 46–50). In our own previous work, we found that the localization of TCRαβ molecules on microvilli is disturbed upon disruption of the actin cytoskeleton with latrunculin A (11). We decided to shed further light on the relationship between the actin cytoskeleton and selective localization of TCRαβ molecules.

We first tested the effect of activation-induced T cell spreading on ICAM-1 on the topographical distribution of TCRαβ molecules. TCR stimulation induces strong actin cytoskeleton remodeling and increased adhesion *via* LFA1 integrin and its ligand ICAM1, leading to stable T cell spreading (3, 51, 52). Spreading may influence the structure of microvilli, as shown in chemokine-activated T cells (53), and may in turn affect the TCR distribution. To probe the effect of spreading, we stimulated Jurkat T-cell with anti-CD3 antibodies at 37°C, and at the same time let them interact with an ICAM-1-coated surface for 3 minutes. We fixed the cells, labelled the membrane with FM143fx, and imaged the TCR distribution as described above. CD3-stimulated Jurkat cells readily spread over the ICAM-1 surface (SM, Fig. S3A) and the typical microvilli structure of these cells prior to stimulation was almost entirely lost with essentially all TCRαβ molecules evenly distributed throughout the membrane (SM, Fig. S3B-C).

To observe actin organization inside the microvilli of T cells, we labelled the membrane of human effector T cells and Jurkat T cells with FM143FX and filamentous (F-) actin with Alexa-647 conjugated phalloidin (a selective marker of F-actin, Fig. 5). Using the combination of VA-TIRFM and SLN, we found that actin filaments fill the space within the microvilli in both human T-effector cells and Jurkat cells (Fig. 5A-B). To further confirm these findings, we performed transmission electron microscopy on thin sections of resin-embedded Jurkat cells. We observed parallel actin filaments within microvilli (Fig. 5C-D), similar to the observation of Majstoravich et al. with lymphoma cells (24). Interestingly, these images do not show actin within the cytoplasm of the cells, which is due to the limited penetration depth of TIRF-based microscopy, as has been noted before (46, 51, 54–56). In contrast, we found that a 3D SLN setup can image the actin meshwork deep inside resting Jurkat cells (Fig. 5I-J).

**Fig. 5:**
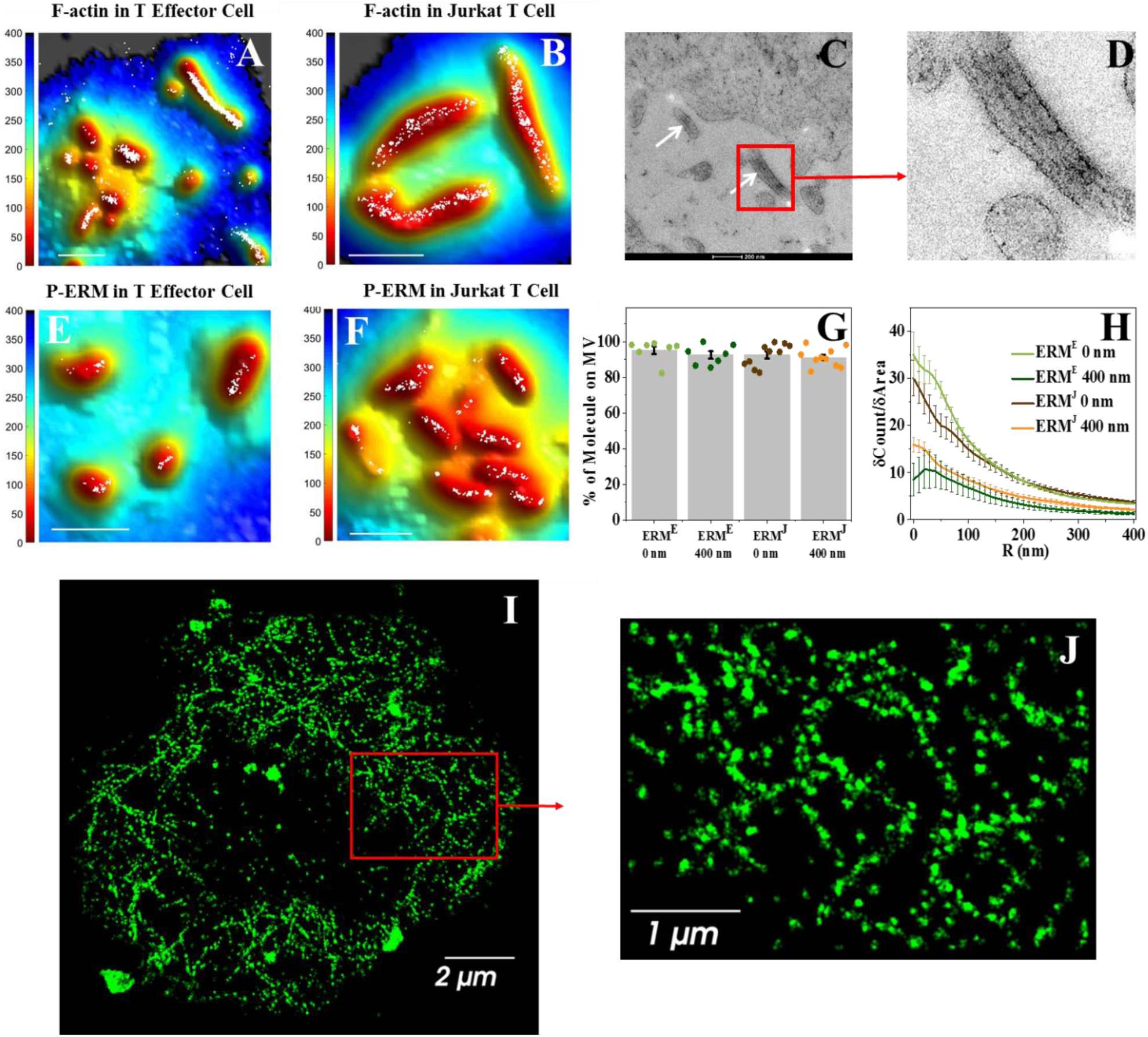
F-Actin and p-ERM both localize within the microvilli of T cells. (A-B) Super-resolution localization maps of Alexa-647-phalloidin labelled F-actin (white dots) overlaid with the 3D surface reconstruction map, showing that F-actin localizes within microvilli. The color bars represent distance from the glass in nm. Scale bars: 1 μm. (C) TEM images of Jurkat cells showing the parallel actin filaments within microvilli (white arrows). (D) The zoomed image of the section indicated by red box in C. (E-G) Super-resolution localization maps of p-ERM molecules (white dots) overlaid with the 3D surface reconstruction map. The color bars represent distance from the glass in nm. Scale bar: 1 μm. (G) Percentage of p-ERM molecules on microvillar (MV) regions at the 0 nm and −400 nm planes for effector cells (ERM^E^) and Jurkat cells (ERM^J^). The values for individual cells are shown as points in the plot. (H) Cumulative increase of the fraction of total molecules on each cell as a function of the distance from the central microvilli region, normalized by the cumulative increase in the fraction of area as a function of distance from microvilli (δCount/δArea). The plots are averages over all cells measured. Error bars represent standard errors of the mean. (I-J) Super-resolution image of the actin meshwork in the cytoplasmic region of a Jurkat cell. (I) Super-resolution image of the actin meshwork in a 1 μm section of the cytoplasmic region of a Jurkat cell, captured by a 3D super-resolution microscope. (J) A zoomed image of the section indicated by a red box in A

ERM (ezrin, radixin and moesin) proteins are well-known for linking the actin cytoskeleton cortex with plasma membrane proteins (57, 58). They also play a crucial role in microvilli formation (53, 59). In T cells only ezrin and moesin are expressed (60). The N-terminal domains (so-called FERM domains (61)) of ERM proteins interact with membrane proteins while their C-terminal domains bind cortical actin (61). Notably, this membrane-cytoskeleton linkage occurs mainly when the ERM C-terminal domain is phosphorylated (62). It has also been shown that after antigen sensing by the TCR, the phosphorylated ERM proteins (referred to as p-ERM) are dephosphorylated, and rapidly de-anchor the actin cytoskeleton cortex from the plasma membrane (63). It was further reported that ERM proteins also control the pre-organization of B cell receptor (BCR) micro-clusters on the plasma membrane of resting B cells (64). Using TEM, Bretscher and coworkers showed early on that ezrin is localized in the apical microvilli of epithelial cells (65). These accumulated findings directed us to investigate the involvement of p-ERM proteins in the anchoring of TCR proteins to microvilli.

We first examined whether p-ERM proteins are localized within microvilli. We labelled p-ERM molecules with Alexa-647 conjugated antibodies and the membrane with FM143FX dye. Combined VA-TIRFM and SLN imaging demonstrated that p-ERM proteins were segregated within microvilli of both human T-effector cells and Jurkat cells (Fig. 5E-F). More than 90% of p-ERM proteins were found to be segregated to the microvillar regions in both imaging planes (Fig. 5G) and the steepness of δCount/δArea plot (Fig. 5H) was very high, supporting the confinement of these molecules within microvilli.

We next reasoned that if the p-ERM proteins act as a bridge to connect TCRαβ to the cortical actin cytoskeleton, these three molecules should co-localize. We therefore performed two-color super-resolution microscopic experiments to find the degree of co-localization of these molecules in pairs on Jurkat T cells with nanometer resolution. We developed a new analysis scheme to obtain the fraction of molecules that are co-localized, described in detail in the Methods section. Our analysis took into account both localization errors and the size of the proteins involved. We first studied co-localization of F-actin and p-ERM (61, 66) by labelling F-actin with alexa-647 conjugated phalloidin and p-ERM with Alexa-568 conjugated anti phospho-ERM antibody and imaging them sequentially by SLN. Although the labelling densities of the two proteins were quite different, we were able to quantify their co-localization (Fig 6A-C). To this end we defined a ‘co-localization fraction’, which is the percentage of molecules of one type that have a molecule of the other type within 45 nm (for details see Methods). The average co-localization fraction, calculated from 10 cells, was 93±2 %. In contrast, the co-localization fraction obtained for other pairs of proteins enriched within microvilli without interacting with each other (L-selectin and CD3, SM, Fig. S4) was only 69±4%. A Kolmogorov-Smirnov test confirmed that the set of co-localization fraction values of p-ERM and F-actin, calculated from individual cell data, differed significantly from the set obtained with L-selectin and CD3, indicating that p-ERM and F-actin closely interact with each other on microvilli.

**Fig. 6:**
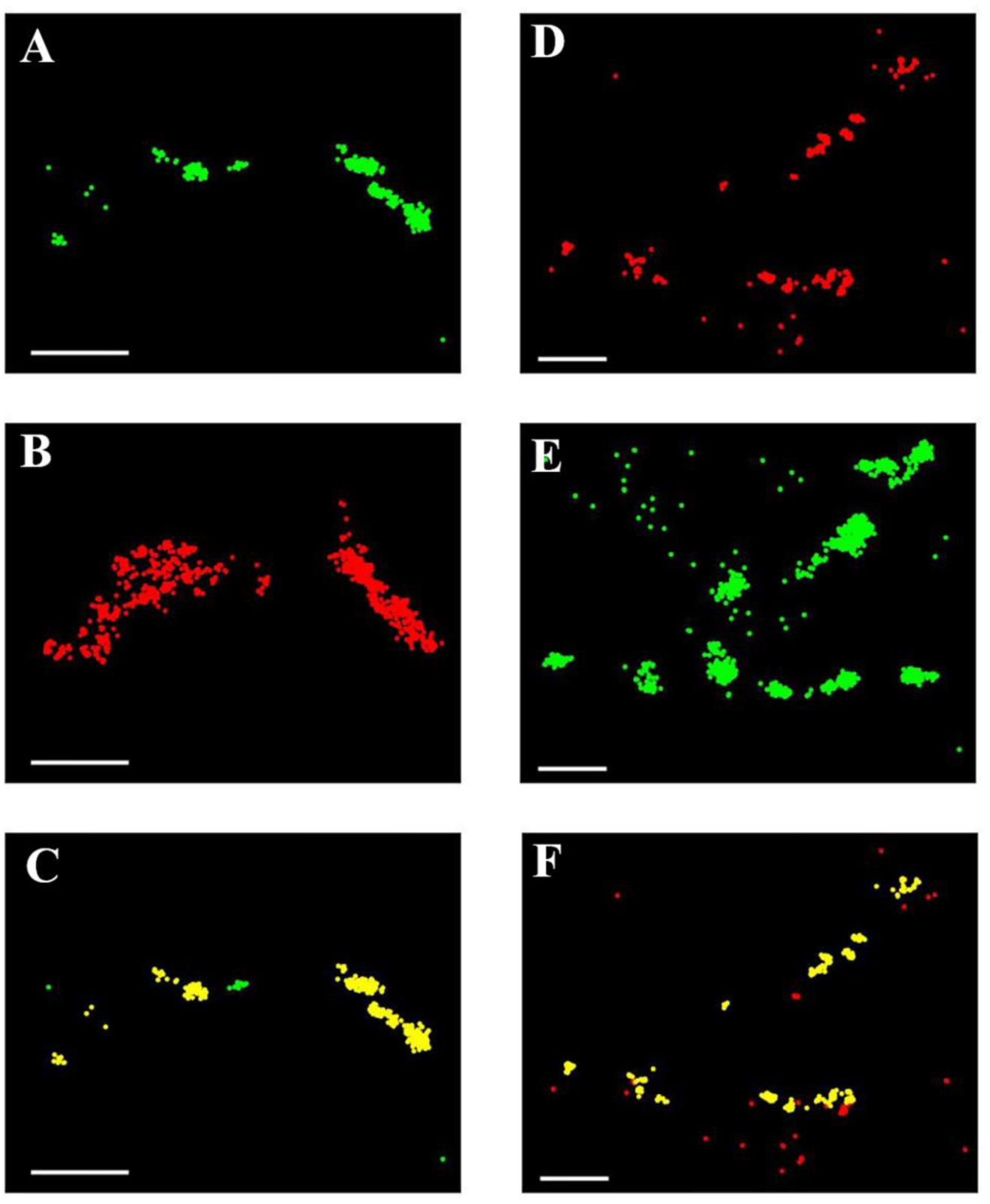
Super-resolution microscopy reveals strong co-localization of p-ERM with F-actin and with TCRαβ. (A) Super-resolution image of a Jurkat cell labelled with Alexa 568 conjugated anti-phospho-ERM antibodies (Green). (B) Super-resolution image of the same cell labelled with Alexa-647 phalloidin (Red), which stains F-actin. (C) Same as A, but p-ERM molecules that have at least one actin molecule within a radius of 45 nm are marked yellow. (D) Super-resolution image of a Jurkat cell labelled with Alexa Fluor 647 anti-human TCRαβ antibodies (Red). (E) Super-resolution image of the same cell labelled with Alexa 568 conjugated anti-phospho-ERM antibodies (Green). (F) Same as in D, but TCRαβ molecules that have at least one p-ERM molecule within a radius of 45 nm are marked yellow. Scale bars: 0.5 μm.

We next followed the co-localization of p-ERM and TCRαβ molecules by labelling p-ERM with Alexa-568 conjugated anti phospho-ERM antibodies as above, and TCRαβ with Alexa-647 conjugated anti-TCRαβ antibodies. The super-resolution images obtained from sequential imaging of the two antibodies showed a high degree of co-localization of TCRαβ and p-ERM (Fig. 6D-F). The average value of the co-localization fraction was 88±1%, which again suggested a high level of interaction between the two proteins, also confirmed by a Kolmogorov-Smirnov test. Taken together, the two co-localization experiments indicated that the TCRαβ heterodimer, p-ERM and F-actin are interacting with each other, suggesting a key role for p-ERM in connecting the TCRαβ complex to the actin cytoskeleton of T cell microvilli.

### Modulating the microvillar distribution of TCRαβ by overexpression of a dominant-negative ezrin fragment

To test the direct role of p-ERM proteins in regulating the formation of the microvilli of T cells, and in maintaining TCRαβ on these microvilli, we transfected T cells with dominant-negative constructs of ezrin. Such constructs have been used as a tool to study the function of ERM proteins in various physiological processes, including TCR-mediated T-cell activation (10, 53, 61, 63). In particular, constructs containing only the N-terminal FERM domain of ezrin and lacking the C-terminal actin-binding domain were shown to block F-actin association with multiple members of the ERM family (7, 53, 61, 63, 67, 68). We transfected Jurkat cells with cDNA encoding a VSV-tagged construct (10) containing the FERM domain of ezrin (which we termed N-ter ezrin) and also with the cDNA encoding of a VSV-tagged construct containing only the C-terminal end of ezrin (C-ter ezrin). N-ter ezrin may only bind with membrane proteins but not with F-actin. It has been reported that N-ter ezrin also causes de-phosphorylation of endogenous p-ERM (63). In contrast, C-ter ezrin cannot bind with membrane proteins as it lacks the FERM domain.

To follow the distribution of TCRαβ distribution on the surface of N-ter ezrin and C-ter ezrin transfected Jurkat cells, we labelled the membrane with FM143FX, TCRαβ with Alexa-647 tagged antibodies and the VSV tag of N-ter ezrin or C-ter ezrin with Alexa-405 conjugated anti-VSV-G epitope tag antibody. Based on the intensity of Alexa-405, we were able to characterize cells as transfected at a high, medium or low level (SM, Fig. S5A-H). Deformation of membrane structure and destruction of microvillar structure correlated with transfection level (SM, Fig. S5I-M). This finding agrees well with Brown et al., who used SEM imaging to show that high N-ter ezrin transfection abolishes microvilli on the surface of Jurkat cells (53). In contrast, the effects of C-ter ezrin transfection on microvilli number and length were negligible (SM, Figs. S5J).

While C-ter ezrin did not affect the localization of TCRαβ to microvilli even at high transfection levels, the distribution of TCRαβ was modified upon increasing N-ter ezrin transfection levels (SM, Fig. S5N-U). The percentage of TCRαβ molecules on the microvilli of C-ter ezrin transfected Jurkat cells was found to be 92.0±1.9%, which is similar to that of non-transfected cells (Fig. 7A), and the δCount/δArea plot for these cells was as steep as that of non-transfected Jurkat cells (Fig. 7B). On the other hand, the percentage of TCRαβ molecules on the microvilli of N-ter ezrin transfected Jurkat cells decreased from 81.0±2.2% for cells with low transfection levels to 59.2±3.8% for cells with medium transfection levels and down to 31.5±3.6% for cells with high transfection levels (Fig. 7A). The slopes of δCount/δArea plots also decreased in a similar fashion (Fig. 7B). Furthermore, the maximum cluster size of TCRαβ molecules, as obtained from the Ripley’s K function analysis also decreased upon increasing N-ter ezrin transfection efficiency (Fig. 7C). Importantly, the degree of localization of TCRαβ on microvilli was found to be inversely correlated with the level of N-ter ezrin transfection (Fig. 7D). This negative correlation confirms our hypothesis that p-ERM proteins maintain the TCRαβ complex on the microvilli of T cells.

**Figure 7.**
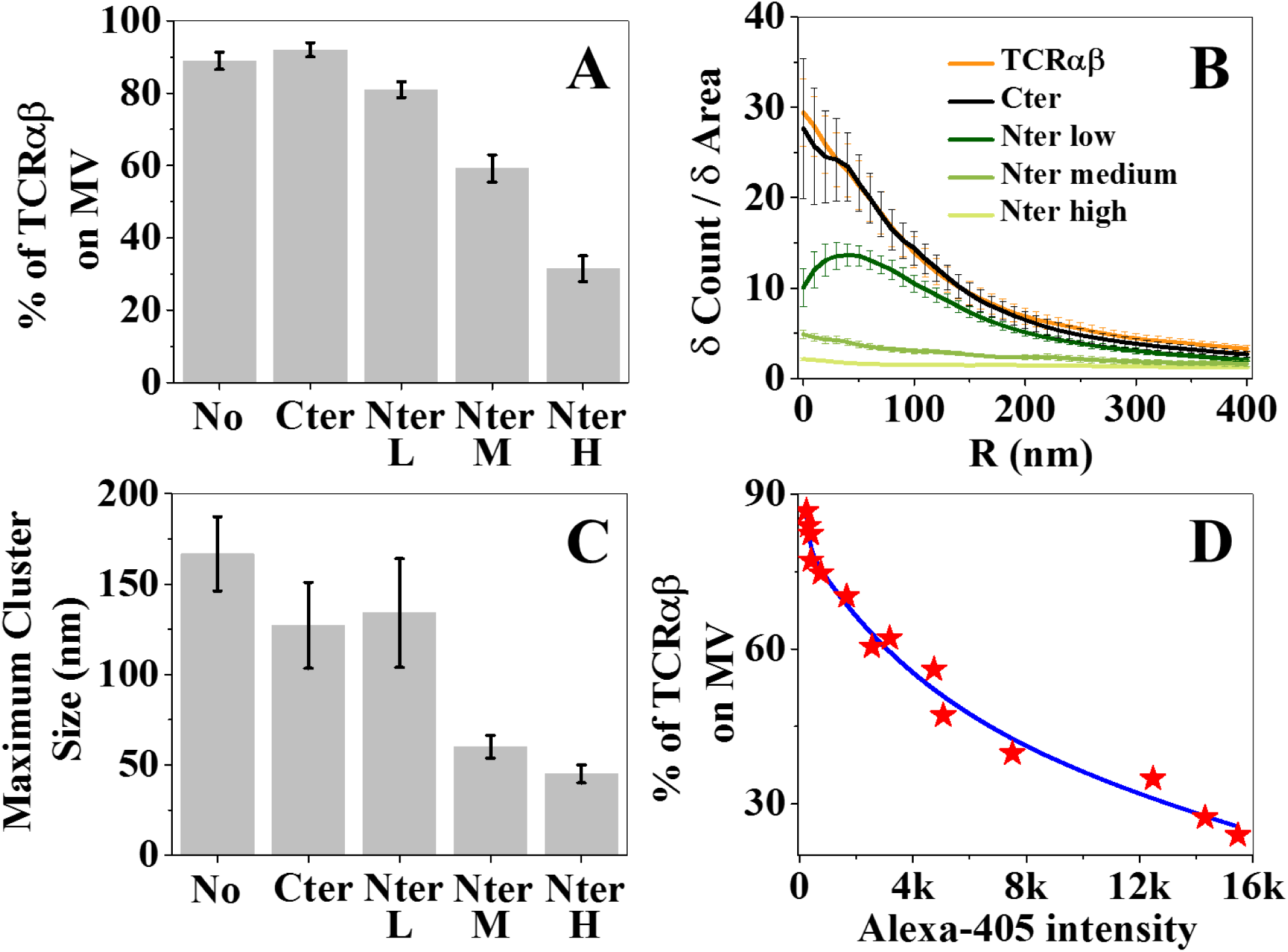
Effect of dominant-negative ezrin transfection on the localization of TCRαβ with respect to 3D topography of Jurkat T cells. (A) Percentage of TCRαβ molecules on microvilli as a function of different transfection levels. ‘No’ refers to no transfection, ‘L’-low, ‘M’-medium, ‘H’-high. (B) Cumulative increase of the fraction of total molecules on each cell as a function of the distance from the central microvilli region, normalized by the cumulative increase in the fraction of area (δCount/δArea plot). The plots are averages over all cells measured. Error bars represent standard errors of the mean. (C) Change of maximum cluster sizes of TCRαβ molecules in response to different transfection levels. Labels as in A. (D) Correlation of percentage of TCRαβ molecules in the microvilli regions and the transfection efficiency, as measured through Alexa-405 intensity. The blue line is a guide to the eye.

We then addressed the functional consequences of altered distribution of TCRs in relation to microvilli. To this end, Jurkat cells transfected with N-ter ezrin were stimulated by incubating them with Raji cells loaded with the superantigen staphylococcal enterotoxin E (SEE), or Raji cells alone as a control (13). We then quantified ERK1/ERK2 kinase activation as a measure of TCR-induced signaling by monitoring phosphorylation levels using flow cytometry with intracellular staining. As shown in SM, Fig. S6A (left panel), we gated on cells expressing either high or low levels of the N-ter ezrin construct (N-ter ezrin-negative cells were not included in the analysis since they contain a mixture of both Jurkat and Raji cells). Interestingly, the percentage of phospho-ERK (pERK) positive cells was significantly higher in cells expressing low N-ter ezrin than in those with high N-ter ezrin expression (SM, Fig. S6A, right panel). Based on three independent experiments, the average percentage of pERK-positive cells was 22.7±2% in low N-ter ezrin expressing cells, and only 12.6±2.5% in high N-ter ezrin expressing cells (SM, Fig. S6B). This difference was found significant with p<0.001. Our results indicate that higher expression of N-ter ezrin, leading to TCR displacement from microvilli, results in impaired T cell activation.

## DISCUSSION

The T-cell surface is covered by numerous elastic actin-rich finger-like protrusions, known as microvilli. Although T cell microvilli have been classically implicated in lymphocyte adhesion to blood vessels (69), the role of these membrane structures in T-cell recognition of antigen presenting cells inside various tissues is poorly understood. Multiple models that attempt to relate T-cell function with the organization of receptors on the cell surface treat the plasma membrane as a 2-D sheet, rather than a 3-D surface (14–18, 20, 48). Recent studies have begun to unearth the pivotal role of microvilli as sensors involved in the initial stages of the immune response (11, 12, 27, 28). In this work we investigated the properties of microvilli that makes them effective sensors.

To that end, we probed the localization of multiple T-cell membrane proteins with respect to microvilli. We mapped the positions of seven key membrane molecules, known to be involved in the initial stages of the immune response, with respect to the 3D topography of the T-cell membrane, both in human effector cells and Jurkat T cells. We found that many (most?) of these proteins reside almost exclusively on microvilli. Among these are components of the TCR complex, including TCRαβ and TCRζ. (Another component of the TCR complex, CD3ε, was demonstrated to reside on microvilli in our previous work (11).) Importantly, CD4, the co-receptor that engages the MHC-II-peptide complex and improves TCR sensitivity by increasing its proximity to its proximal tyrosine kinase Lck (40), is also co-localized with its TCR partner on microvilli. Furthermore, the co-stimulatory adhesion molecule CD2 was also found to be segregated on microvilli. This indicates that in addition to the TCR and its co-receptor complex, critical co-stimulatory receptors are also enriched on microvilli, probably in proximity with the TCR complexes, thus amplifying the TCR signals delivered during the initial stages of T cell activation. Like CD2, CD28 also functions as a co-stimulatory receptor that amplifies TCR signals in naïve T cells (70). It would be therefore interesting to investigate in future work whether CD28 is also enriched on microvilli, and whether CD2 and CD28 co-localize on these cellular projections. Two proteins critically involved in the initial TCR signaling steps, namely Lck and LAT, were also found to be enriched on microvilli, with ∼76% and ∼68% of these molecules, respectively, located on these T cell projections. Thus, all the major TCR signaling molecules necessary for the initial immune response are pre-organized together on T cell microvilli. It remains unknown, however, why a significant fraction of Lck and LAT molecules are found on cell body regions rather than on microvilli. Interestingly, a pool of Lck molecules that are not associated with CD4 was reported by Stephen et al. (41) It is therefore possible that this Lck pool resides on the cell body of T cells and operates on a different subset of substrates. Moreover, some Lck and LAT molecules were found in endosomal vesicles, whose position with respect to the plasma membrane may vary (71, 72).

A recent report (73) suggested that observations of TCR clusters on the T-cell membrane (17, 18, 20, 22, 23, 48) were due to imaging artifacts, and that all TCR machineries are randomly distributed on the surface of T cells. However, in that paper TCR-stimulated T cells were spread on ICAM-1-coated surfaces. Our own experiments with ICAM-1-coated surfaces conclusively indicate that the observation of random distribution of TCRs on the plasma membrane of T cell is likely the outcome of both the TCR stimulation and the spreading process on ICAM-1. In contrast to the T cells studied on ICAM-1, which model APC-bound T cells, our studies of the distribution of various surface molecules on microvilli involved resting T cells fixed prior to APC encounter. These T cells retained their preformed morphology as well as the preformed segregation of their TCR machineries and proximal signaling molecules on their microvilli. It should be noted that the clustering of T-cell proteins we observed here has a clear mechanistic origin, i.e. segregation on microvilli. Thus, our results link for the first time the preorganization of TCR assemblies on resting T cells to the presence of microvilli, which are constantly scanning for antigenic signals on various APCs (27).

To shed light on the mechanism of selective anchoring of TCR associated molecules on microvilli, we studied the distribution of TCRαβ receptors with respect to actin filaments and their phosphorylated ERM linkers. The co-localization of these three components, and the negative effects of a DN-ezrin on TCRαβ-actin assemblies, collectively suggest that the actin cytoskeleton is intimately involved in the organization of TCR assemblies within microvilli via its ERM linkers (either directly or indirectly) in the organization of the preformed TCR assemblies within microvilli. ERM-dependent actin anchoring may form a chemical and a physical barrier for the diffusion of TCR molecules away from microvilli. This type of a barrier might not completely prevent lateral diffusion of TCR molecules. Indeed, single-particle tracking experiments on TCR components demonstrated lateral motion, but found diffusion coefficients (74, 75) that are ∼100 times smaller than the typical diffusion coefficients of free membrane proteins (76).

The connection of TCR components to the actin filaments inside microvilli also allows T cells to exert significant forces on these components during the first few seconds following encounter of the APC, and these forces may stabilize the TCR-MHC-peptide bonds, as suggested by recent biophysical measurements(77–80). As the T cell interaction with the APC persists, the phosphorylated ERM linkers become inactivated by dephosphorylation(63) and release the connection of the cortical actin cytoskeleton and the plasma membrane in the vicinity of the TCR machineries. The TCRαβ and its proximal signaling molecules likely use multiple ERM proteins as switchable phosphorylation-regulated linkers, which diffuse away from the tips of the microvilli they are located on following TCR activation and dephosphorylation.

Taken together, our results demonstrate that the pre-organization of the TCR machinery on microvilli depends on the intact anchorage to the actin cytoskeleton facilitated by ERM linkers. The pre-organization of TCR complexes and co-stimulatory molecules on T-cell microvilli may have several advantages. First, the flexible structure and constant motion of microvilli likely make the search for antigens on APCs faster and more efficient (27). Indeed, recent work emphasized the importance of the initial encounter between the T cell and the APC and the fast signal generation that includes conformational changes in relevant proteins such as Lck (81), followed by phosphorylation of downstream kinases like ZAP-70 as well as calcium mobilization (4–6). Second, the large surface area of microvilli accommodates a large number of membrane proteins. Third, the concentration of TCR molecules with their proximal signaling molecules on microvilli increases their avidity for cognate antigenic peptides on the APC and thereby facilitates TCR signal transmission. Indeed, recent reports highlight the ability of individual TCR microclusters to deliver local signals within seconds or a few minutes, (27, 78, 82, 83) prior to the formation of stable ISs that consist of large TCR supramolecular structures (84).

In conclusion, we demonstrate that rather than being randomly distributed on the T-cell membrane, multiple proteins involved in the initial recognition and incorporation of antigenic signals are specifically pre-clustered on microvilli, which facilitates the initial recognition events underlying the TCR activation cascade. Upon recognition of cognate peptide/MHC complexes on APCs, one or more of these TCR machineries likely get released from their original assemblies within microvilli, and together with gradual collapse of the microvilli and additional remodeling of the actin cytoskeleton (85, 86) promote the formation of stable immune synapses. Future work should focus on this putative TCR release process, and help identify if and how dephosphorylation of ERM family members facilitates the redistribution of TCR machineries and the topographical alterations that enable them to drive optimal T cell activation.

## Methods

### Key Resources Table

| REAGENT or RESOURCE | SOURCE | IDENTIFIER |
| --- | --- | --- |
| **Antibody** |  |  |
| Alexa 647-conjugated mouse anti-human TCR$\alpha\beta$ antibody | Biolegend (clone IP26) | Cat# 306714 |
| Purified anti-human TCR$\alpha\beta$ antibody | Biolegend (clone IP26) | Cat# 306702 |
| Alexa 647-conjugated mouse anti-human CD45 antibody | Biolegend (clone HI30) | Cat# 304020 |
| Alexa 647-conjugated anti-human CD62L antibody | Biolegend (clone DREG-56) | Cat# 304818 |
| Alexa 647 anti-human CD3 antibody | Biolegend (clone HIT3a) | Cat# 300322 |
| Purified anti-human CD3 antibody | Biolegend (clone OKT3) | Cat# 317302 |
| Purified anti-CD247 (other names: TCRζ, CD3ζ) antibody | Biolegend (clone 6B10.2) | Cat# 644102 |
| Purified anti-LAT antibody | Biolegend (clone LAT1111) | Cat# 623902 |
| Purified anti-Lck antibody | Biolegend (clone LCK-01) | Cat# 628302 |
| Purified anti-human CD2 antibody | Biolegend (clone RPA-2.10) | Cat# 300202 |
| Purified anti-human CD4 antibody | Biolegend (clone OKT4) | Cat# 317402 |
| Purified anti-VSV-G epitope tag antibody | Biolegend (clone Poly29039) | Cat# 903901 |
| Anti-phospho-ezrin (Thr567)/radixin (Thr564)/moesin (Thr558) (48G2) rabbit antibody | CST Technology | mAb #3726 |
| **F-actin probe** |  |  |
| Alexa Fluor™ 647 phalloidin | Thermo Fisher Scientific | Cat# A22287 |
| **Dyes** |  |  |
| Alexa Fluor™ 405 | Thermo Fisher Scientific | Cat# A30000 |
| Alexa Fluor™ 568 | Thermo Fisher Scientific | Cat# A20103 |
| Alexa Fluor™ 647 | Thermo Fisher Scientific | Cat# A20106 |
| **Software** |  |  |
| MATLAB | MathWorks | N/A |
| Origin | OriginLab Corporation | N/A |

### Experimental Model and Subject Details

#### Human T cells

To isolate human peripheral T lymphocytes, de-identified whole blood from healthy donors was citrate-anticoagulated, followed by dextran sedimentation and Ficoll (Sigma) gradient separation. To remove B cells, a nylon wool column (Unisorb) was used. After incubating the cells in a complete RPMI growth medium for >2 h, only the non-adherent fraction of cells was collected to a new plate (this step resulted in ∼90% CD3+ T lymphocytes). To generate effector T cells, the cells were stimulated on an anti-CD3/anti-CD28 antibody-coated plate for 2 d. After that, the effector cells were cultured in a complete RPMI growth medium that contained IL-2 (350 U/mL) and 2-mercaptoethanol (50 μM) for 7–14 d.

#### Jurkat T cell culture

Jurkat clone E6.1 cells were grown in RPMI media containing 10% fetal bovine serum (FBS), 10 mM Hepes, 100 unit/ml penicillin, 100 µg/ml streptomycin and 2 mM L-glutamine in an atmosphere of 5% (v/v) CO2 enriched air at 37 °C.

### Method Details

#### Transfection

Jurkat E6.1 cells were washed in PBS without Ca^2+^ and Mg^2+^. They were then re-suspended in Neon resuspension buffer R (ThermoFisher Scientific) at 1×10^8^ cells/ml, and mixed with plasmid DNA (20 µg/sample) expressing either the N-terminal or the C-terminal ezrin domain tagged with a VSV-G epitope (a kind gift from Dr. M. Arpin (87)). 100 µl cells were electroporated with three 1400 V pulses, 10 ms each, using a Neon transfection system (ThermoFisher Scientific). Cells were transferred to a RPMI-1640 medium with Glutamax-I medium (Gibco/ThermoFisher Scientific), supplemented with 10% fetal bovine serum without antibiotics and cultured overnight at +37°C in 5% CO2.

#### ERK1/ERK2 kinase activation assay

Jurkat cells were transfected either with an empty vector or with the N-ter ezrin expression plasmid as described above. The day after, cells were collected, washed and mixed with Raji cells, either untreated or pre-loaded with bacterial superantigen (staphylococcal enterotoxin E, SEE, Toxin Technology, Inc.; 15 µg/ml;). After incubation for 10 min at 37°C, cells were fixed in 4% paraformaldehyde for 15 min, washed twice in PBS then permeabilized with ice-cold methanol and left overnight at −20°C. Cells were then washed twice in PBS + 0.5% BSA (FACS buffer) and stained with an anti-VSV-G monoclonal antibody (clone P5D4, purified from ascites), to assess the expression level of the N-ter ezrin, and anti-pERK1/2 rabbit antibodies (Cell Signaling Technology; ref. #9101). Samples were then washed twice in the FACS buffer and incubated with anti-mouse IgG1-PE (BD Biosciences, ref.#550083) and anti-rabbit IgG-AlexaFluor647 (ThermoFisher Scientific, ref. #A21245) conjugates. Finally, cells were washed, resuspended in the FACS buffer and analyzed with a MacsQuant flow cytometer (Miltenyi Biotech). Data analysis was performed using FlowJo 10 (FlowJo, LLC). Statistical significance was assessed with a one-way ANOVA test using GraphPad Prism software.

#### Antibody labeling

All unlabeled antibodies were labeled with Alexa Fluor™ 405, Alexa Fluor™ 568, Alexa Fluor™ 647 according to need. For this, unlabeled antibody in PBS buffer was reacted with the NHS ester of the corresponding dye in a 1:10 ratio in presence of 0.1 M sodium bicarbonate buffer for 1 h at room temperature in the dark. Micro Bio-Spin column with Bio-Gel P-30 (Bio-Rad) was used to remove the unlabeled dye molecules.

#### Cellular labeling with antibodies

A suspension of Jurkat or effector T cells was washed with 5 mM EDTA/PBS for 5 min by centrifugation at 4 °C. The cells were incubated in a blocking solution (1% BSA, 5 mM EDTA, 0.05% N_3_Na, PBS) on ice for 10 min. For labeling of surface molecules, the cells were treated with antibodies (10-20 μg/mL according to the antibody) for 20-60 min on ice. After washing the cells twice with 5 mM EDTA/PBS by centrifugation at 4 °C, they were fixed with a fixation buffer [4 % (wt/vol) paraformaldehyde, 0.2–0.5% glutaraldehyde, 2% (wt/vol) sucrose, 10 mM EGTA, and 1 mM EDTA, PBS] in suspension for 2 h on ice. The fixative was washed twice with PBS by centrifugation. The cell membrane was then stained with FM143FX (Invitrogen; 5 μg/mL) for 30 min on ice. Another fixation for 30 min on ice was then performed and fixatives were washed twice at 4 °C with PBS by centrifugation. Cells were suspended in PBS and were kept at 4 °C. These cells were then directly used for super-resolution mapping of the receptors/adhesion molecule of interest. We verified that the procedure above, involving fixation in solution at 4 °C, captures the true resting state of the cells-see Supporting Text.

For intracellular staining of signaling proteins, cells were first fixed with a fixation buffer [4 % (wt/vol) paraformaldehyde, 0.2–0.5% glutaraldehyde, 2% (wt/vol) sucrose, 10 mM EGTA, and 1 mM EDTA, PBS] in suspension for 2 h on ice. The fixative was washed twice with PBS by centrifugation. The cell membrane was then stained with FM1-43FX for 30 min on ice. Another fixation for 30 min on ice was performed and fixatives were washed twice at 4 °C with PBS by centrifugation. Cells were then incubated in a permeabilization buffer (0.05% saponin and 1 % BSA in PBS) along with the antibody (10 μg/mL) at 4 °C. Incubation time varied according to the antibody. After that, cells were washed twice at 4°C with PBS by centrifugation. Cells were suspended in PBS and were kept at 4 °C. To rule out alteration of the morphology of Jurkat T cells by these procedures, we imaged fixed cells with SEM (Fig. 1A-B). We found that the microvilli-dominated surface structure of Jurkat cells was intact.

For actin labelling, Jurkat cells were washed with the cytoskeleton buffer (50 mM imidazole, 50 mM KCl, 0.5 mM MgCl2, 0.1 mM EDTA, and 1 mM EGTA (pH 6.8)), incubated for 30 min at room temperature and then fixed with a fixation buffer [4 % (wt/vol) paraformaldehyde, 0.2–0.5% glutaraldehyde in cytoskeleton buffer] in suspension for 2 h on ice. After washing the cells with PBS, the cell membrane was labeled with FM1-43FX for 30 min on ice. Cells were then incubated in a permeabilization buffer (0.05% saponin and 1 % BSA in PBS) along with the Alexa Fluor™ 647 phalloidin (1 unit/mL) at 4°C. They were washed twice at 4°C with PBS by centrifugation and immediately imaged.

For the single molecule tracking study of TCRαβ, Jurkat cells were labeled with Atto 647N conjugated mouse anti-human TCRαβ antibodies (3 μg/mL) for 20 min on ice. The cells were washed twice with 5 mM EDTA/PBS and then used live.

For dual-color super resolution microscopy, two different proteins were labeled sequentially, using the same protocol described above.

For imaging studies with dominant negative ERM transfected Jurkat cells, TCRαβ was labeled on live cells that were then fixed. The membrane was labeled with FM1-43FX and cells were fixed again using the same protocol described above. The cells were then incubated in a permeabilization buffer (0.05% saponin and 1 % BSA in PBS) along with the Alexa Fluor™ 405 labeled anti-VSV-G epitope tag antibody (10 μg/mL) at 4 °C. The cells were washed twice at 4 °C with PBS by centrifugation, suspended in PBS and kept at 4 °C.

#### Sample preparation for microscopy

Glass-bottom petri dishes were cleaned with 1 M NaOH (Fluka) for 40 min, then coated with PLL (0.01%; Sigma) for 40 min and washed with PBS. 100 µl of the labeled cell solution in PBS was placed in the petri dish and cells were allowed to settle on the PLL surface for 10 minutes. The PBS buffer was then exchanged with a freshly prepared “blinking buffer” [50 mM cysteamine (Sigma), 0.5 mg/mL glucose oxidase (Sigma), 40 μg/mL catalase (Sigma), 10% (wt/vol) glucose (Sigma), 93 mM Tris·HCl, PBS buffer (pH 7.5–8.5)] and the sample was kept for 30 minutes before imaging was conducted.

#### TIRF setup

Our microscope setup was described in detail in our previous publication (11). Briefly, the setup had three different lasers sources (405 nm; Toptica iBEAM-SMART-405-S, 532 nm; COBOLT SambaTM 50 mW and 642 nm; Toptica iBEAM-SMART-640-S). A series of dichroic beam splitters (z408bcm, z532bcm; Chroma) and an objective lens (M-20×; N.A., 0.4; Newport) were used to couple the lasers into a polarization-maintaining single-mode fiber. This fiber was connected to an acousto-optic tunable filter (AOTFFnc-VIS-TN-FI; AA Opto-Electronic), which was used to separately modulate each excitation beam. A polarizer (GT10-A; Thorlabs) for each laser was used to modulate the polarization to maximize coupling efficiency. We controlled the power of each laser either manually by adjustment of a half-wave plate (WPH05M; Thorlabs) positioned after a polarizer or by the computer. Achromatic lenses (01LAO773, 01LAO779; CVI Melles Griot) were utilized to expand and collimate the first-order output beams from the AOTF to a diameter of 6 mm. An achromatic focusing lens (f = 500 mm;LAO801; CVI Melles Griot) was employed to focus the expanded laser beams at the back focal plane of the microscope objective lens (UAPON 100×OTIRF; N.A., 1.49; Olympus). To achieve total internal reflection at the sample we shifted the position of the focused beam from the center of the objective to its edge. A quad-edge super-resolution laser dichroic beam splitter (Di03-R405/488/532/635-t1-25×36) was utilized to separate fluorescence emitted from the excitation. It was then coupled out from the side port of the microscope (Olympus IX71). Notch filters (NF01-405/488/532/635 StopLine Quad-notch filter and ZET635NF; Semrock) were used to block the residual scattered laser light. A single EMCCD camera (iXonEM +897 back-illuminated; Andor) was employed in a dual-view mode: Spectrally separated images were projected onto the two halves of the CCD chip. Our setup had a final magnification of 240×, which resulted in a pixel size of 66.67 nm.

#### Reconstruction of 3D cell surfaces and detection of microvilli

The methodology for 3D surface reconstruction and detection of microvilli from TIRF images taken at a series of angles was described in detail in reference (11). Briefly, we recorded TIRF images of cells at a series of angles of incidence under weak illumination of a 532-nm laser. The pixel with maximum intensity at a particular angle of incidence (*I*_max_(θ)) should be the closest one to the glass surface. To calculate the relative axial distance (*δz*) of each point on the cell surface we used the following equation;

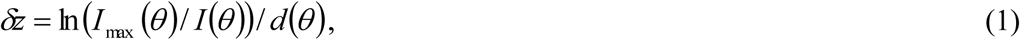

where *d* (*θ*) represents penetration depth of the TIRF evanescent field with an angle of incidence θ, which may be calculated from the following equation:

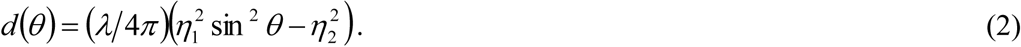

In this equation λ stands for the excitation wavelength (532 nm), and η_1_ and η_2_ represent the refractive index of the immersion oil (1.52) and blinking buffer (1.35), respectively.

To find the location of the tips of microvilli, we developed an algorithm in MATLAB. First, the “Laplacian of a Gaussian” (LoG) filter was applied to the TIRF images, and the processed images were then converted to binary images by setting a threshold. The binary images were segmented using the function “bwlabel”. To include the maximum number of microvilli, the binary image acquired from the data recorded at the smallest angle of incidence was considered for segmentation first, and then data from the image recorded at the largest angle of incidence was combined to separate individual microvilli. Finally, segmented areas obtained from images taken at angles 65.5°, 66.8°, and 68.2° were combined and the mean of the δz of each pixel was calculated and used to generate the LocTips map (Fig. S1).

#### SLN measurement and drift correction

Alexa-647 and Alexa-568 dyes were used for SLN measurements. These dyes can be converted to a dark state upon illumination with a very high intensity laser of the appropriate wavelength. In the presence of a buffer containing β-mercaptoethylamine together with glucose oxidase/catalase (an oxygen-scavenging system (34, 88)), a small fraction of the dye molecules may be switched back at any moment of time. Thus, individual molecules appear separately in consecutive recording frame. This phenomenon helps to detect the location of a molecule with precision well below the diffraction limit. For all SLN measurements, the laser beam incident angle was 66.8°. For each focal plane, we recorded 28,000 frames, in 7 series of movies (each containing 4000 frames). We used a piezo stage (PI nano Z-Piezo slide scanner; PI) to record SLN movies at the 0 nm and −400 nm focal planes. The measurement at two different planes ensured that proteins localized on membrane regions situated somewhat further from the surface due to the length of microvillar protrusions would not be overlooked. In fact, given the significant depth of focus, and given the high laser power used in the super-resolution measurements, focusing on the −400 nm plane allowed us to better observe cell body regions, while still obtaining signals from molecules on microvilli as well.

For drift correction, we recorded TIRF reference images of the cell membrane between SLN movies under weak illumination of a 532-nm laser (10–20 μW). We used 2D cross-correlation of each reference TIRF image with the last reference image taken in a series for identifying the drift level. For dual-color SLN, we used fluorescent nanodiamonds (a kind gift of Dr. Keir Neumann, NIH, Bethesda, MD) as fiducial points for drift correction.

#### Localization analysis

Detailed molecular localization procedures were provided in reference (1). Briefly, a thresholding step was used to identify individual emitters. They were then fitted to a 2D Gaussian function to obtain their x and y coordinates and the widths of their point-spread functions in the x and y directions. We obtained a narrow distribution of widths in these measurements. Only those molecules whose widths were within ±3 SD of the mean were considered. The average localization uncertainty in x–y was estimated to be ∼11 nm (FWHM), while the image resolution, calculated based on Fourier ring correlation analysis of images (89), was ∼30 nm. (Note that the latter depends both on the localization uncertainty and on the density of labeled molecules.)

#### Quantitative analysis of molecular distribution

To analyze the distribution of each membrane protein, we segmented the membrane area of each cell into microvilli (MV) regions and non-microvilli or cell-body (CB) regions using the analysis described in our previous work (11). We then calculated percentage of molecules on microvilli regions and on cell body regions (SM, Fig. S7A-D) of each cell.

We also determined the cumulative fractional increase of the number of molecules of a specific protein on each cell as a function of the distance from the central region of each microvillus (SM, Fig. S7E-G). To this end, the central region of each microvillus was defined as the region that is not more than 20 nm from the microvillar tip, which is the pixel the minimum δz value. We then used the ‘boundary’ function of MATLAB to encapsulate this central region (SM, Fig. S7F), which could take different shapes, depending on the shape of the specific microvillus and how it sat with respect to the surface. We then plotted concentric closed curves of a similar shape and increasing size (SM, Fig. S7G), and calculated the number of molecules in each concentric structure, from which we obtained the cumulative fractional increase. This value was normalized by the cumulative fractional increase of the area, to obtain the δCount/δArea plot. The more a protein localized to the microvillar region, the steeper was the slope of this plot.

#### Cluster Analysis

To estimate clustering of membrane proteins, we used Ripley’s K-function (18, 90), which calculates the number of molecules within distance r of a randomly chosen molecule:

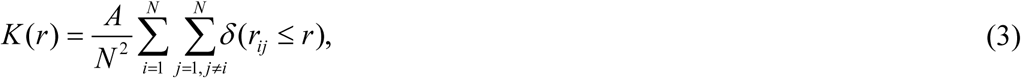

where A represents the image area of interest, N is the total number of localizations in the area of interest, and *δ* (*r_ij_* ≤ *r*) equals 1 if the distance between the two molecules (*r_ij_*) is smaller or equal to r. Molecules were only included if they were at least at a distance r away from the edge of the image. We used a linear transformation of the K-function to get an estimate of maximum cluster sizes as follows:

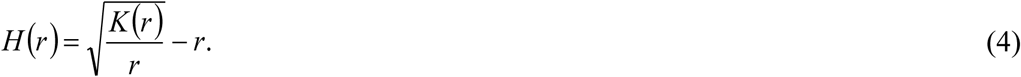

For a random distribution of particles, H(r) equals zero. Positive values of H(r) represent clustering of particles. The maximum amplitude of H(r) represents the size of the largest clusters (90). To verify our cluster analysis protocol, we simulated clusters with sizes normally distributed around a known value. We then performed the cluster analysis on the simulated data. The maximum cluster size calculated from the cluster analysis on the simulated data agreed well with the known maximum cluster size.

#### Quantitative estimation of the co-localization level of membrane proteins

We developed a simple but effective method for estimating co-localization of proteins, which relied on measuring distances between localized points in super resolved images. In particular, we defined the co-localization percentage of molecules i and j as 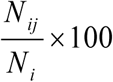, where *N_i_* is the total number of detected points of molecule i, and *N_ij_* is the number of points of molecule i that have at least one point of molecule j within the interaction distance (defined below). For the molecule i we always selected the molecule with the lower labeling density.

To specify the interaction distance for two proteins, one needs to take into account two factors: the image resolution (∼30 nm in our case, calculated based on Fourier ring correlation analysis of images (89)) and the maximum distance of two proteins during interaction. Interestingly, the P-ERM proteins can elongate up to 30 nm when bound with actin filaments (91). Therefore, we took our interaction distance as ∼45 nm (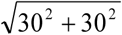).

We evaluated the super-resolution co-localization percentage obtained under a scenario in which there is no interaction between molecules. We considered the case of two molecules that co-exist on microvilli without interaction. For this purpose, we performed dual-color localization experiments of L-selectin and CD3 (See SM, Fig. S4 for details). The average value of co-localization percentage for these two molecules was ∼ 68%. An average co-localization percentage significantly higher than 68% would therefore indicate considerable interaction between two molecules.

#### 3D SLN setup

Three-dimensional super resolution images of the actin cytoskeleton were collected on a Vutara SR352 (Bruker) microscope based on the single-molecule localization biplane technology, using a 60x silicon oil immersion objective (1.3 NA). Alexa Fluor 647 phalloidin labeled actin was excited using 640 nm laser (5 kW/cm2), and 45,000 frames were acquired with an acquisition time of 20 ms per frame. Imaging was performed in the presence of the imaging buffer (7 μM glucose oxidase (Sigma), 56 nM catalase (Sigma), 5 mM cysteamine (Sigma), 50 mM Tris, 10 mM NaCl, 10% glucose, pH 8). Data were analyzed and visualized by the Vutara SRX software.

#### SEM Imaging

Please refer to our previous publication for detailed procedures (11). Briefly, we fixed Jurkat cells using 4% (wt/vol) paraformaldehyde, 0.5%glutaraldehyde, 2% (wt/vol) sucrose, 10 mM EGTA, and 1 mM EDTA in PBS for 2 h on ice. We placed the fixed cells on a PLL coated coverslip. Different concentrations of ethanol were used to dehydrate the cells. Then the samples were dried in a BAL-TEC CPD 030 critical point drier. We placed the samples on a carbon tape and coated with carbon (Edwards; Auto306) or sputter-coated with gold–palladium (Edwards; S150). A Ultra 55 SEM (Zeiss) was used to image the samples.

#### TEM imaging

Sample preparation was adapted from Eltsov et al. (92) Cells were concentrated by centrifugation. A drop of the pellet was placed on an aluminum disc (Engineering Office M. Wohlwend GmbH, Sennwald, Switzerland) with a 100 µm deep cavity and covered with a flat disc. The sample was then frozen in a HPM 010 high-pressure freezing machine (Bal-Tec, Liechtenstein). Cells were subsequently freeze substituted (AFS2, Leica Microsystems, Vienna, Austria) in anhydrous acetone containing 0.5% uranyl acetate (prepared from 20% stock in absolute methanol). After incubation for 42 hours at −90 °c, the temperature was raised during 24 h to −30 °c and the samples were washed 3 times in anhydrous acetone, infiltrated with increasing concentrations of Lowicryl HM20 resin (4h 10%, 4h 30%, overnight 60%, 4h 90%, 4h 100%, overnight 100%, 4h 100%) and polymerized under ultraviolet light. Thin sections (70 nm) were examined in a Tecnai Spirit T12 electron microscope (FEI, Eindhoven, the Netherlands) operating at 120 kV. Images were recorded with an Eagle 2k × 2k CCD camera (FEI).

## Acknowledgements

We thank Drs. Sandeep Yadav, Alexander Vaskevich and Yunmin Jung, as well as Mr. Francesco Roncato of the Weizmann Institute of Science for their kind advice and experimental help during the project. We also thank Dr. Keir Neuman (NIH) for kindly sharing with us fluorescent nanodiamond samples. The SEM studies were conducted at the Electron Microscopy Unit of the Weizmann Institute of Science. G.H. is the incumbent of the Hilda Pomeraniec Memorial Professorial Chair.

## References

1. Dustin ML (2014) What counts in the immunological synapse? Mol Cell 54(2):255–262.

2. Monks CR, Freiberg BA, Kupfer H, Sciaky N, & Kupfer A (1998) Three-dimensional segregation of supramolecular activation clusters in T cells. Nature 395(6697):82–86.

3. Grakoui A, et al. (1999) The immunological synapse: a molecular machine controlling T cell activation. Science 285(5425):221–227.

4. Henrickson SE, et al. (2013) Antigen availability determines CD8(+) T cell-dendritic cell interaction kinetics and memory fate decisions. Immunity 39(3):496–507.

5. Marangoni F, et al. (2013) The transcription factor NFAT exhibits signal memory during serial T cell interactions with antigen-presenting cells. Immunity 38(2):237–249.

6. Friedl P & Brocker EB (2002) TCR triggering on the move: diversity of T-cell interactions with antigen-presenting cells. Immunol Rev 186:83–89.

7. Malissen B & Bongrand P (2015) Early T cell activation: integrating biochemical, structural, and biophysical cues. Annu Rev Immunol 33:539–561.

8. Kaizuka Y, Douglass AD, Vardhana S, Dustin ML, & Vale RD (2009) The coreceptor CD2 uses plasma membrane microdomains to transduce signals in T cells. J Cell Biol 185(3):521–534.

9. Klammt C & Lillemeier BF (2012) How membrane structures control T cell signaling. Front Immunol 3:291.

10. Roumier A, et al. (2001) The membrane-microfilament linker ezrin is involved in the formation of the immunological synapse and in T cell activation. Immunity 15(5):715–728.

11. Jung Y, et al. (2016) Three-dimensional localization of T-cell receptors in relation to microvilli using a combination of superresolution microscopies. Proc Natl Acad Sci U S A 113(40):E5916–E5924.

12. Razvag Y, Neve-Oz Y, Sajman J, Reches M, & Sherman E (2018) Nanoscale kinetic segregation of TCR and CD45 in engaged microvilli facilitates early T cell activation. Nat Commun 9(1):732.

13. Lasserre R, et al. (2010) Ezrin tunes T-cell activation by controlling Dlg1 and microtubule positioning at the immunological synapse. EMBO J 29(14):2301–2314.

14. Davis SJ & van der Merwe PA (2006) The kinetic-segregation model: TCR triggering and beyond. Nat Immunol 7(8):803–809.

15. Sezgin E, Levental I, Mayor S, & Eggeling C (2017) The mystery of membrane organization: composition, regulation and roles of lipid rafts. Nat Rev Mol Cell Biol 18(6):361–374.

16. Simons K & Sampaio JL (2011) Membrane organization and lipid rafts. Cold Spring Harb Perspect Biol 3(10):a004697.

17. Lillemeier BF, et al. (2010) TCR and Lat are expressed on separate protein islands on T cell membranes and concatenate during activation. Nat Immunol 11(1):90–96.

18. Roh KH, Lillemeier BF, Wang F, & Davis MM (2015) The coreceptor CD4 is expressed in distinct nanoclusters and does not colocalize with T-cell receptor and active protein tyrosine kinase p56lck. Proc Natl Acad Sci U S A 112(13):E1604–1613.

19. Ritter AT, et al. (2017) Cortical actin recovery at the immunological synapse leads to termination of lytic granule secretion in cytotoxic T lymphocytes. Proc Natl Acad Sci U S A 114(32):E6585–E6594.

20. Sherman E, et al. (2011) Functional nanoscale organization of signaling molecules downstream of the T cell antigen receptor. Immunity 35(5):705–720.

21. Rossy J, Owen DM, Williamson DJ, Yang Z, & Gaus K (2013) Conformational states of the kinase Lck regulate clustering in early T cell signaling. Nat Immunol 14(1):82–89.

22. Pageon SV, et al. (2016) Functional role of T-cell receptor nanoclusters in signal initiation and antigen discrimination. Proc Natl Acad Sci U S A 113(37):E5454–5463.

23. Hu YS, Cang H, & Lillemeier BF (2016) Superresolution imaging reveals nanometer- and micrometer-scale spatial distributions of T-cell receptors in lymph nodes. Proc Natl Acad Sci U S A 113(26):7201–7206.

24. Majstoravich S, et al. (2004) Lymphocyte microvilli are dynamic, actin-dependent structures that do not require Wiskott-Aldrich syndrome protein (WASp) for their morphology. Blood 104(5):1396–1403.

25. Nijhara R, et al. (2004) Rac1 mediates collapse of microvilli on chemokine-activated T lymphocytes. J Immunol 173(8):4985–4993.

26. Schwarz US & Alon R (2004) L-selectin-mediated leukocyte tethering in shear flow is controlled by multiple contacts and cytoskeletal anchorage facilitating fast rebinding events. Proc Natl Acad Sci U S A 101(18):6940–6945.

27. Cai E, et al. (2017) Visualizing dynamic microvillar search and stabilization during ligand detection by T cells. Science 356(6338).

28. Kim HR, et al. (2018) T cell microvilli constitute immunological synaptosomes that carry messages to antigen-presenting cells. Nat Commun 9(1):3630.

29. Goodfellow HS, et al. (2015) The catalytic activity of the kinase ZAP-70 mediates basal signaling and negative feedback of the T cell receptor pathway. Sci Signal 8(377):ra49.

30. Stock K, et al. (2003) Variable-angle total internal reflection fluorescence microscopy (VA-TIRFM): realization and application of a compact illumination device. J Microsc 211(Pt 1):19–29.

31. Sundd P, et al. (2010) Quantitative dynamic footprinting microscopy reveals mechanisms of neutrophil rolling. Nat Methods 7(10):821–824.

32. Rust MJ, Bates M, & Zhuang X (2006) Sub-diffraction-limit imaging by stochastic optical reconstruction microscopy (STORM). Nat Methods 3(10):793–795.

33. Endesfelder U & Heilemann M (2015) Direct stochastic optical reconstruction microscopy (dSTORM). Methods Mol Biol 1251:263–276.

34. van de Linde S, et al. (2011) Direct stochastic optical reconstruction microscopy with standard fluorescent probes. Nat Protoc 6(7):991–1009.

35. Alcover A, Alarcon B, & Di Bartolo V (2018) Cell Biology of T Cell Receptor Expression and Regulation. Annu Rev Immunol 36:103–125.

36. Taniuchi I (2018) CD4 Helper and CD8 Cytotoxic T Cell Differentiation. Annu Rev Immunol 36:579–601.

37. Jardine L, et al. (2013) Rapid detection of dendritic cell and monocyte disorders using CD4 as a lineage marker of the human peripheral blood antigen-presenting cell compartment. Front Immunol 4:495.

38. Roose JP, et al. (2003) T cell receptor-independent basal signaling via Erk and Abl kinases suppresses RAG gene expression. PLoS Biol 1(2):E53.

39. Rossy J, Williamson DJ, & Gaus K (2012) How does the kinase Lck phosphorylate the T cell receptor? Spatial organization as a regulatory mechanism. Front Immunol 3:167.

40. Foti M, Phelouzat MA, Holm A, Rasmusson BJ, & Carpentier JL (2002) p56Lck anchors CD4 to distinct microdomains on microvilli. Proc Natl Acad Sci U S A 99(4):2008–2013.

41. Stephen TL, Wilson BS, & Laufer TM (2012) Subcellular distribution of Lck during CD4 T-cell maturation in the thymic medulla regulates the T-cell activation threshold. Proc Natl Acad Sci U S A 109(19):7415–7420.

42. Williamson DJ, et al. (2011) Pre-existing clusters of the adaptor Lat do not participate in early T cell signaling events. Nat Immunol 12(7):655–662.

43. Makgoba MW, Sanders ME, & Shaw S (1989) The CD2-LFA-3 and LFA-1-ICAM pathways: relevance to T-cell recognition. Immunol Today 10(12):417–422.

44. Wilkins AL, Yang W, & Yang JJ (2003) Structural biology of the cell adhesion protein CD2: from molecular recognition to protein folding and design. Curr Protein Pept Sci 4(5):367–373.

45. Fernandes RA, et al. (2019) A cell topography-based mechanism for ligand discrimination by the T cell receptor. Proc Natl Acad Sci U S A 116(28):14002–14010.

46. Fritzsche M, et al. (2017) Cytoskeletal actin dynamics shape a ramifying actin network underpinning immunological synapse formation. Sci Adv 3(6):e1603032.

47. Burkhardt JK, Carrizosa E, & Shaffer MH (2008) The actin cytoskeleton in T cell activation. Annu Rev Immunol 26:233–259.

48. Lillemeier BF, Pfeiffer JR, Surviladze Z, Wilson BS, & Davis MM (2006) Plasma membrane-associated proteins are clustered into islands attached to the cytoskeleton. Proc Natl Acad Sci U S A 103(50):18992–18997.

49. Tamzalit F, et al. (2019) Interfacial actin protrusions mechanically enhance killing by cytotoxic T cells. Sci Immunol 4(33).

50. Roybal KT, et al. (2015) Modest Interference with Actin Dynamics in Primary T Cell Activation by Antigen Presenting Cells Preferentially Affects Lamellal Signaling. PLoS One 10(8):e0133231.

51. Bunnell SC, Kapoor V, Trible RP, Zhang W, & Samelson LE (2001) Dynamic actin polymerization drives T cell receptor-induced spreading: a role for the signal transduction adaptor LAT. Immunity 14(3):315–329.

52. Stewart MP, Cabanas C, & Hogg N (1996) T cell adhesion to intercellular adhesion molecule-1 (ICAM-1) is controlled by cell spreading and the activation of integrin LFA-1. J Immunol 156(5):1810–1817.

53. Brown MJ, et al. (2003) Chemokine stimulation of human peripheral blood T lymphocytes induces rapid dephosphorylation of ERM proteins, which facilitates loss of microvilli and polarization. Blood 102(12):3890–3899.

54. Smoligovets AA, Smith AW, Wu HJ, Petit RS, & Groves JT (2012) Characterization of dynamic actin associations with T-cell receptor microclusters in primary T cells. J Cell Sci 125(Pt 3):735–742.

55. Babich A, et al. (2012) F-actin polymerization and retrograde flow drive sustained PLCgamma1 signaling during T cell activation. J Cell Biol 197(6):775–787.

56. Bunnell SC, et al. (2002) T cell receptor ligation induces the formation of dynamically regulated signaling assemblies. J Cell Biol 158(7):1263–1275.

57. Fehon RG, McClatchey AI, & Bretscher A (2010) Organizing the cell cortex: the role of ERM proteins. Nat Rev Mol Cell Biol 11(4):276–287.

58. McClatchey AI (2014) ERM proteins at a glance. J Cell Sci 127(Pt 15):3199–3204.

59. Tsukita S & Yonemura S (1999) Cortical actin organization: lessons from ERM (ezrin/radixin/moesin) proteins. J Biol Chem 274(49):34507–34510.

60. Shaffer MH, et al. (2009) Ezrin and moesin function together to promote T cell activation. J Immunol 182(2):1021–1032.

61. Bretscher A, Edwards K, & Fehon RG (2002) ERM proteins and merlin: integrators at the cell cortex. Nat Rev Mol Cell Biol 3(8):586–599.

62. Gautreau A, Louvard D, & Arpin M (2000) Morphogenic effects of ezrin require a phosphorylation-induced transition from oligomers to monomers at the plasma membrane. J Cell Biol 150(1):193–203.

63. Faure S, et al. (2004) ERM proteins regulate cytoskeleton relaxation promoting T cell-APC conjugation. Nat Immunol 5(3):272–279.

64. Treanor B, Depoil D, Bruckbauer A, & Batista FD (2011) Dynamic cortical actin remodeling by ERM proteins controls BCR microcluster organization and integrity. J Exp Med 208(5):1055–1068.

65. Berryman M, Franck Z, & Bretscher A (1993) Ezrin is concentrated in the apical microvilli of a wide variety of epithelial cells whereas moesin is found primarily in endothelial cells. J Cell Sci 105 (Pt 4):1025–1043.

66. Arpin M, Chirivino D, Naba A, & Zwaenepoel I (2011) Emerging role for ERM proteins in cell adhesion and migration. Cell Adh Migr 5(2):199–206.

67. Allenspach EJ, et al. (2001) ERM-dependent movement of CD43 defines a novel protein complex distal to the immunological synapse. Immunity 15(5):739–750.

68. Delon J, Kaibuchi K, & Germain RN (2001) Exclusion of CD43 from the immunological synapse is mediated by phosphorylation-regulated relocation of the cytoskeletal adaptor moesin. Immunity 15(5):691–701.

69. von Andrian UH, Hasslen SR, Nelson RD, Erlandsen SL, & Butcher EC (1995) A central role for microvillous receptor presentation in leukocyte adhesion under flow. Cell 82(6):989–999.

70. Esensten JH, Helou YA, Chopra G, Weiss A, & Bluestone JA (2016) CD28 Costimulation: From Mechanism to Therapy. Immunity 44(5):973–988.

71. Bouchet J, et al. (2017) Rab11-FIP3 Regulation of Lck Endosomal Traffic Controls TCR Signal Transduction. J Immunol 198(7):2967–2978.

72. Soares H, et al. (2013) Regulated vesicle fusion generates signaling nanoterritories that control T cell activation at the immunological synapse. J Exp Med 210(11):2415–2433.

73. Rossboth B, et al. (2018) TCRs are randomly distributed on the plasma membrane of resting antigen-experienced T cells. Nat Immunol 19(8):821–827.

74. Dushek O, et al. (2008) Effects of intracellular calcium and actin cytoskeleton on TCR mobility measured by fluorescence recovery. PLoS One 3(12):e3913.

75. Favier B, Burroughs NJ, Wedderburn L, & Valitutti S (2001) TCR dynamics on the surface of living T cells. Int Immunol 13(12):1525–1532.

76. Fujiwara TK, et al. (2016) Confined diffusion of transmembrane proteins and lipids induced by the same actin meshwork lining the plasma membrane. Mol Biol Cell 27(7):1101–1119.

77. Liu Y, et al. (2016) DNA-based nanoparticle tension sensors reveal that T-cell receptors transmit defined pN forces to their antigens for enhanced fidelity. Proc Natl Acad Sci U S A 113(20):5610–5615.

78. Sibener LV, et al. (2018) Isolation of a Structural Mechanism for Uncoupling T Cell Receptor Signaling from Peptide-MHC Binding. Cell 174(3):672–687 e627.

79. Feng Y, et al. (2017) Mechanosensing drives acuity of alphabeta T-cell recognition. Proc Natl Acad Sci U S A 114(39):E8204–E8213.

80. Depoil D & Dustin ML (2014) Force and affinity in ligand discrimination by the TCR. Trends Immunol 35(12):597–603.

81. Philipsen L, et al. (2017) De novo phosphorylation and conformational opening of the tyrosine kinase Lck act in concert to initiate T cell receptor signaling. Sci Signal 10(462).

82. Taylor MJ, Husain K, Gartner ZJ, Mayor S, & Vale RD (2017) A DNA-Based T Cell Receptor Reveals a Role for Receptor Clustering in Ligand Discrimination. Cell 169(1):108–119 e120.

83. Varma R, Campi G, Yokosuka T, Saito T, & Dustin ML (2006) T cell receptor-proximal signals are sustained in peripheral microclusters and terminated in the central supramolecular activation cluster. Immunity 25(1):117–127.

84. Dustin ML & Choudhuri K (2016) Signaling and Polarized Communication Across the T Cell Immunological Synapse. Annu Rev Cell Dev Biol 32:303–325.

85. Hui KL, Balagopalan L, Samelson LE, & Upadhyaya A (2015) Cytoskeletal forces during signaling activation in Jurkat T-cells. Mol Biol Cell 26(4):685–695.

86. Vicente-Manzanares M & Sanchez-Madrid F (2004) Role of the cytoskeleton during leukocyte responses. Nat Rev Immunol 4(2):110–122.

87. Algrain M, Turunen O, Vaheri A, Louvard D, & Arpin M (1993) Ezrin contains cytoskeleton and membrane binding domains accounting for its proposed role as a membrane-cytoskeletal linker. J Cell Biol 120(1):129–139.

88. Dempsey GT, et al. (2009) Photoswitching mechanism of cyanine dyes. J Am Chem Soc 131(51):18192–18193.

89. Nieuwenhuizen RP, et al. (2013) Measuring image resolution in optical nanoscopy. Nat Methods 10(6):557–562.

90. Gao J, et al. (2015) Revealing the cellular localization of STAT1 during the cell cycle by super-resolution imaging. Sci Rep 5:9045.

91. Jayasundar JJ, et al. (2012) Open conformation of ezrin bound to phosphatidylinositol 4,5-bisphosphate and to F-actin revealed by neutron scattering. J Biol Chem 287(44):37119–37133.

92. Eltsov M, et al. (2015) Quantitative analysis of cytoskeletal reorganization during epithelial tissue sealing by large-volume electron tomography. Nat Cell Biol 17(5):605–614.

