## Supplementary Data for "ERM-Dependent Assembly of T-Cell Receptor Signaling and Co-stimulatory Molecules on Microvilli Prior to Activation"

### Supporting Text

**Labelling at 4°C captures the bona fide resting state of T cells.** Cells used in this work were fixed in solution in order to guarantee that the true resting state of the cells is obtained and avoid any activation due to interaction with the surface [3]. However, even in solution cells might be activated due to interaction with some antibodies at physiological temperature (37°C). We therefore elected to perform labelling at 4°C. To verify that the lower temperature does not create any labelling artifacts, we performed the following experiments. First, we the compared staining of T cells with anti Alexa-647 labelled L-selectin and anti-CD45 antibodies at 4°C and 37°C, following fixation at the same temperature in each case (Fig. S1). We found that at both temperatures L-selectin was concentrated on microvilli, [1,2] while CD45 was distributed uniformly throughout the membrane. [1,4,5] Next, we stained T cells with Alexa-647 labelled anti human TCRαβ antibodies at the two temperatures. At 4°C the structure of microvilli on almost all the T cells was intact, and the TCRαβ molecules localized on them (Fig. S1T). On the other hand, at 37°C most of the cells lost their microvillar structures, likely due to stimulation by the antibodies. We did find a few T cells that still presented intact microvillar structure, and in these cells, TCRαβ was localized on the microvilli (Fig. S1U), similar to the observation at 4°C. Thus, it is evident that T-cell labelling and fixation at 4°C in solution helps capturing the bona fide resting state of the cells.

**Molecules in the two planes probed in SLN are detected with equal probability.** One can argue that due to the exponentially decaying intensity of the TIRF laser field, the detection of molecules closer to glass surface might be biased. To check this issue we imaged two different glass surface uniformly coated with Alexa-647 dye molecules (Fig. S1M-O). In the first experiment, the dye molecules were uniformly coated directly on the glass surface. Using the TIRF microscope, we captured videos on at least 10 different regions (8.5 μm X 8.5 μm each) of the glass surface, focusing right at the interface with the glass substrate (Fig. S1M). Using photobleaching steps to count molecules, we found that the average number of molecules detected within a region was ~26.4± 1.5 (Fig. S1O). In the second experiment, the glass surface was first coated with a 500 nm thick layer of the polymer MY-133 MC (MY Polymers, Israel), whose refractive index (1.328) is almost same as that of water. Dye molecules were then coated uniformly on top of the polymer layer. We now captured videos while focusing 500 nm away from the glass substrate. The average number of molecules detected within a region was in this case ~25.8± 2.1 (Fig. S1N-O). Clearly, our dual-plane SLN technique is able to detect molecules even when they are 500 nm away from glass surface. This rules out the possibility of a biased detection of molecules closer to glass surface.

**Microvilli can be identified using L-selectin localization.** We observed elongated bulging structures (0.5-1 μ long) in our TIRF images (Fig 1F). We attributed these structures to microvilli that lie down horizontally on the glass surface. In fact, microvilli that are flattened against the cell body can be observed even in SEM images, as demonstrated in Fig. S1J-K (see red arrows). To further validate the identity of the elongated bulging structures, we carefully studied L-selectin localization relative to these structures. It is well known that L-selectin exclusively localizes on the microvilli of resting lymphocyte cells[1,2], which makes this surface molecule a most reliable marker of microvilli. Labelling Jurkat cells with anti-L-selectin antibodies, we observed that ridges were enriched with L-selectin (black arrow in Fig. S1L), confirming their identity as microvilli lying on their side.

### Supporting Figures

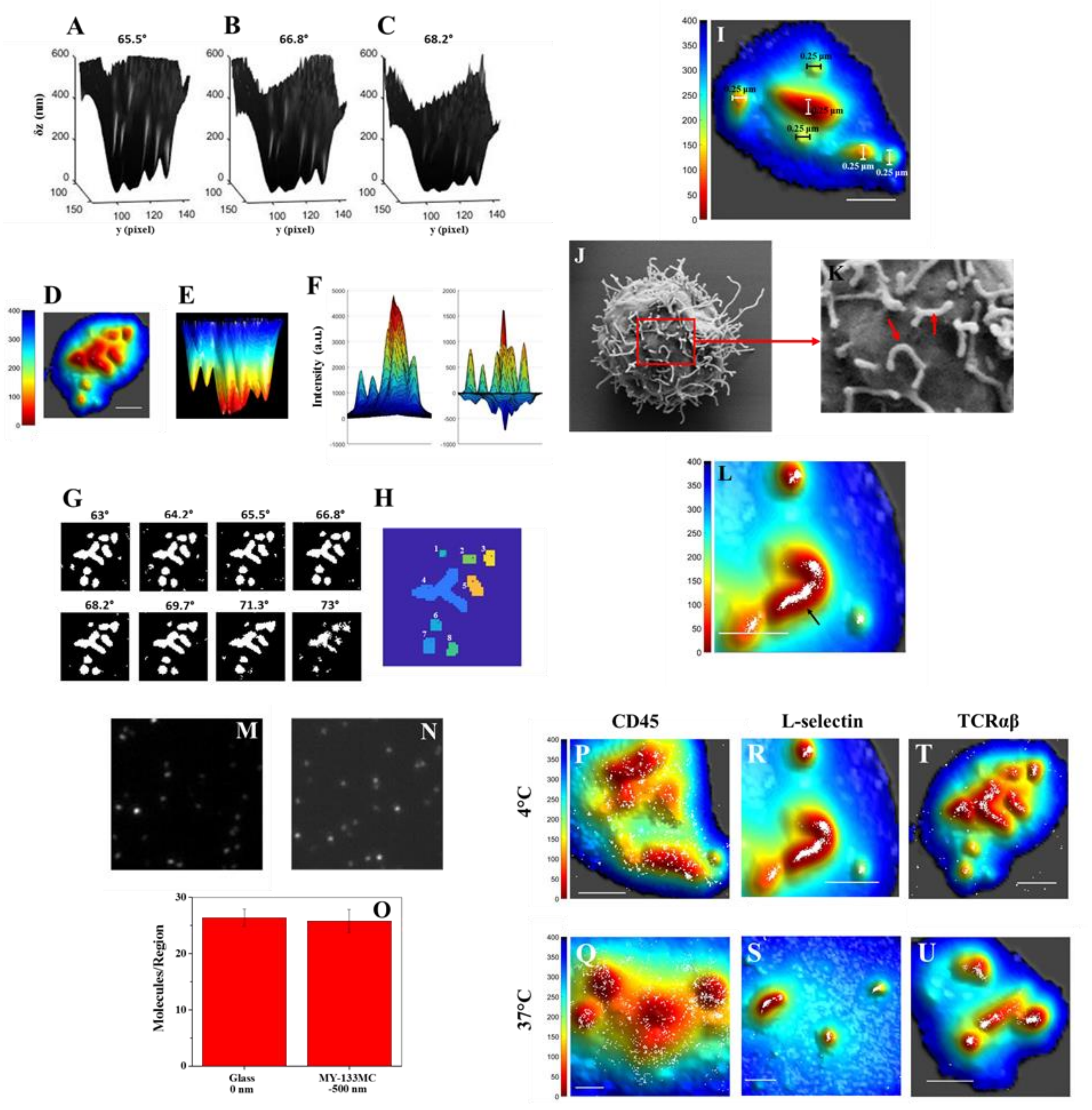

**Figure S1. Validation of Methodology (Related to Figure 1)**

Reconstruction of 3D surface of a T cell from VA-TIRFM and determination of microvilli regions (the LocTips map). (A–C) A series of 3D surface reconstruction maps of a Jurkat T cell from TIRFM images, based on calculated δz values from TIRFM measurements at angles of incidence 65.5° (A), 66.8° (B), and 68.2° (C) under weak illumination of a 532-nm laser. The δz map is shown from a direction perpendicular to the y–z plane. (D) A representative 3D surface reconstruction map of a T cell, calculated as the mean of δz values of A–C. The δz values are represented by different hues with a step size of 6.25 nm. (E) A bottom-to-top 2D projection of D. (Scale bar: 1 µm.) (F) The regions above a threshold value of 5 in Laplacian-of-Gaussian filter images (white) obtained from a series of images acquired at a series of angles of incidence (indicated above each image). (G) The segmentation map of distinguishable protruding areas is determined by combining a series of segmentation analyses of VA-TIRFM measurements at incident angles from 63° to 73°. Each individual microvillus is numbered. The coordinates of the pixel of minimum δz in each individual microvillus are marked (black cross). Using the segmentation maps of multiple cells, we could estimate that the densities of microvilli on the Jurkat and effector T-cell surface are 1.8 ± 0.2 and 3 ± 0.5[1] per $\mu m^2$ respectively. This density compares well with the density obtained from SEM images, which are 2.5 ± 0.1 and 4.5 ± 0.5[1] per $\mu m^2$ respectively, and with literature values (2–4 per $\mu m^2$)[6]

Identification of Microvilli: (I) Widths of microvilli structures are diffraction-limited in 3D surface reconstruction maps. (J) SEM image of a Jurkat T cells and (K) zoomed region of the SEM image demonstrating microvilli lying flat against the cell body (red arrows). (L) Positions of L-selectin molecules obtained from SLN (green dots) at the 0 nm plane are superimposed on membrane topography maps of Jurkat T cells obtained from VA-TIRFM. (Black arrow denotes an L-selectin enriched microvillus lying down horizontally on the glass.)

Detection of Molecules away from glass surface using dual-plane SLN. (M) First frame of a representative movie taken right at the interface with a glass surface uniformly coated with 50 µl of 15 nM of Alexa-647 dye molecules. (N) First frame of a representative movie taken 500 nm away from a MY-133MC coated glass surface (thickness 500 nm), which was then uniformly coated with 50 µl of 15 nM of Alexa-647 dye molecules. (O) Detected average number of molecules per region in the above cases. Error bar represents SEM.

Localization maps of membrane proteins of T-cell labeled at 4°C and 37°C. Positions of protein molecules obtained from SLN (white dots) at the 0 nm plane on Jurkat T cells are superimposed on membrane topography maps obtained from VA-TIRFM. The color bars represent distance from the glass in nm. Scale bars: 1 µm. (P) CD45 at 4°C, (Q) CD45 at 37°C, (R) L-selectin at 4°C, (S) L-selectin at 37°C, (T) TCRαβ at 4°C and (U) TCRαβ at 37°C.

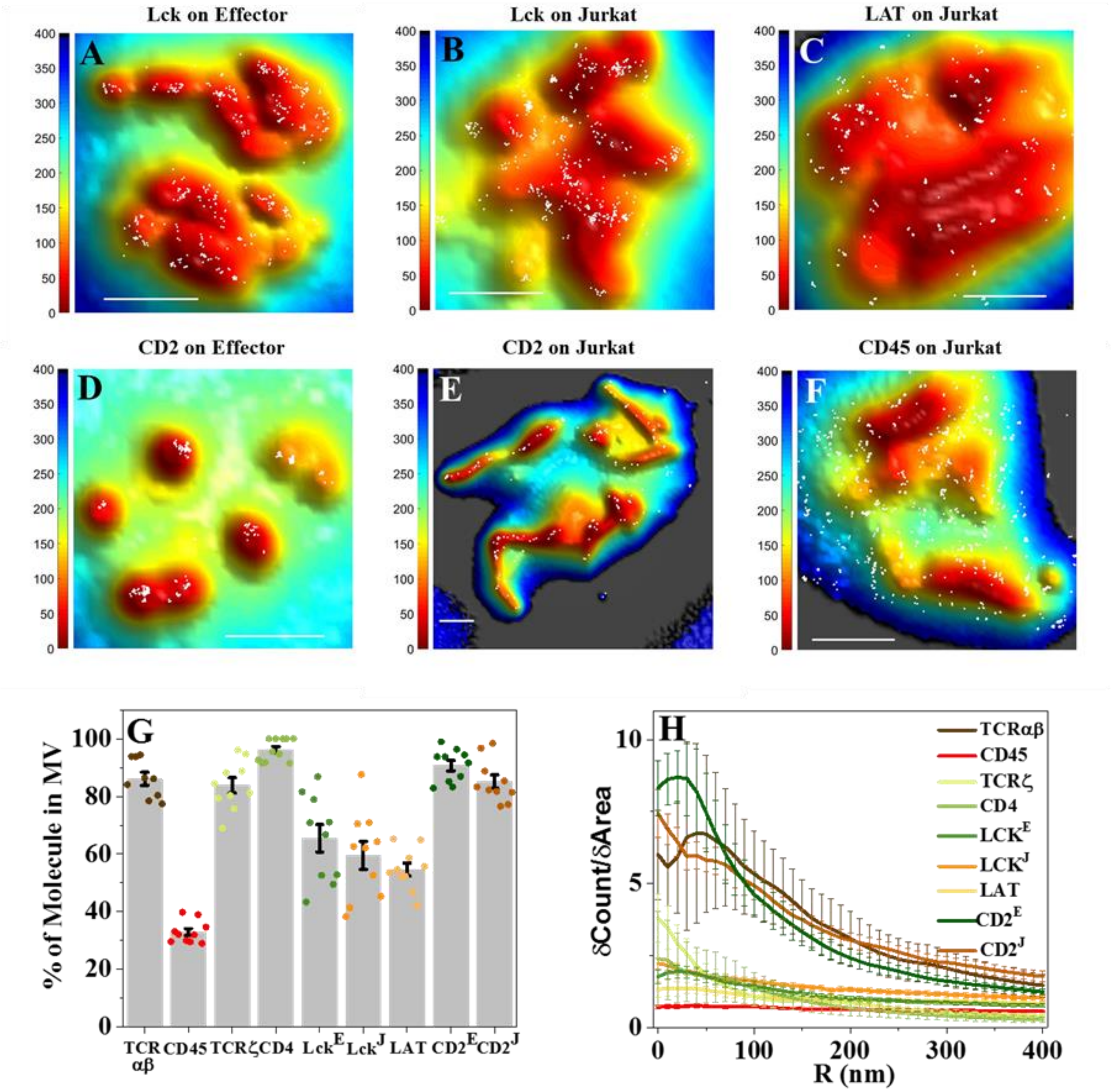


**Figure S2: Localization maps of membrane proteins involved in the initial immune response and quantitative measures for the distribution of receptors on the T cell surface measured at the -400 nm plane (Related to Figs. 2-4).** As in Figure 2, positions of molecules obtained from SLN (white dots) at the -400 nm plane are superimposed on membrane topography maps obtained from VA-TIRFM. The color bars represent distance from glass in nm. Scale bars: 1 μm. The cell type (either Jurkat or human effector) appears in parentheses after the name of the protein. (A) Lck (Effector), (B) Lck (Jurkat), (C) LAT (Jurkat), (D) CD2 (Effector), (E) CD2 (Jurkat) and (F) CD45 (Jurkat).

(G) Percentage of molecules on microvillar (MV) regions of the membrane. The values for individual cells are shown as dots in the plot. (H) Cumulative increase of the fraction of total molecules on each cell as a function of the distance from the central microvilli region, normalized by the cumulative increase in the fraction of area (δCount/δArea) as a function of distance from microvilli. The plots are averages over all cells measured at the -400 nm plane. Error bars represent standard errors of the mean. The E and J superscripts following some protein names denote values obtained with effector cells and Jurkat cells, respectively. When no superscript is shown, the values were obtained with Jurkat cells.

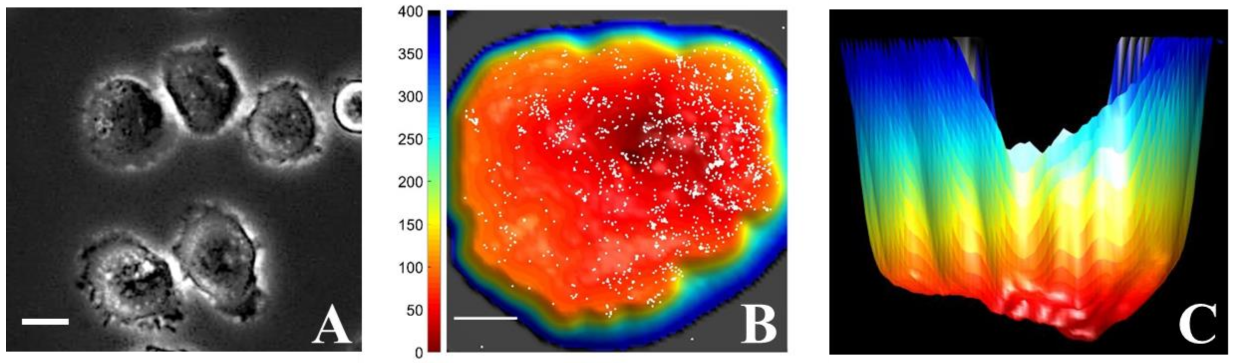


**Figure S3. Effect of CD3-stimulated T-cell spreading on ICAM-1-coated surfaces on the distribution of TCRαβ (Related to Figs. 5-7).** (A) Phase contrast image of OKT3-stimulated (10 μg/ml)) cells interacting with ICAM-1-coated surface. Scale bar 10 μm. (B) TCRαβ molecules are randomly distributed on the surface of the activated and spread cells. White dots are SLN-based localizations superimposed on membrane topography maps obtained from VA-TIRFM. The color bars represent distance from the glass in nm. Scale bar: 1 μm. (C) Side view of the 3D reconstruction map of the same Jurkat cell membrane shown in B.

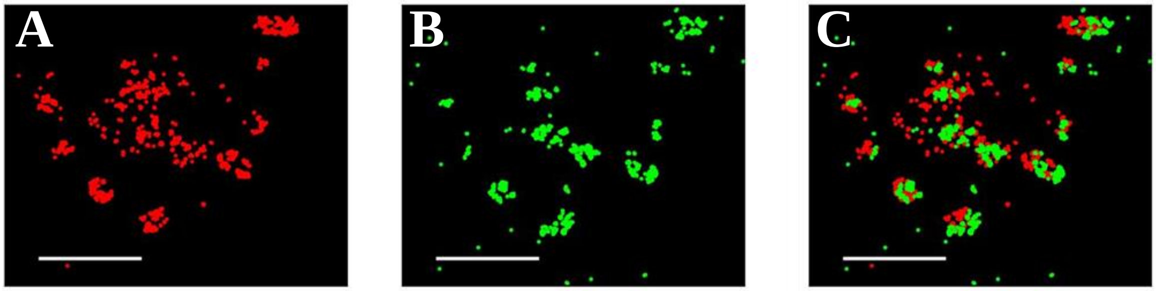


**Figure S4. Super-resolution co-localization of CD3 and L-Selectin on T cell membranes (Related to Fig. 6).** (A) CD3 labelled with an Alexa Fluor 647 conjugated anti-human CD3 antibody (Red). (B) L-Selectin labeled with an Alexa 568 conjugated anti- L-Selectin antibody (Green). (C) A and B superimposed. Scale bar: 1 μm.

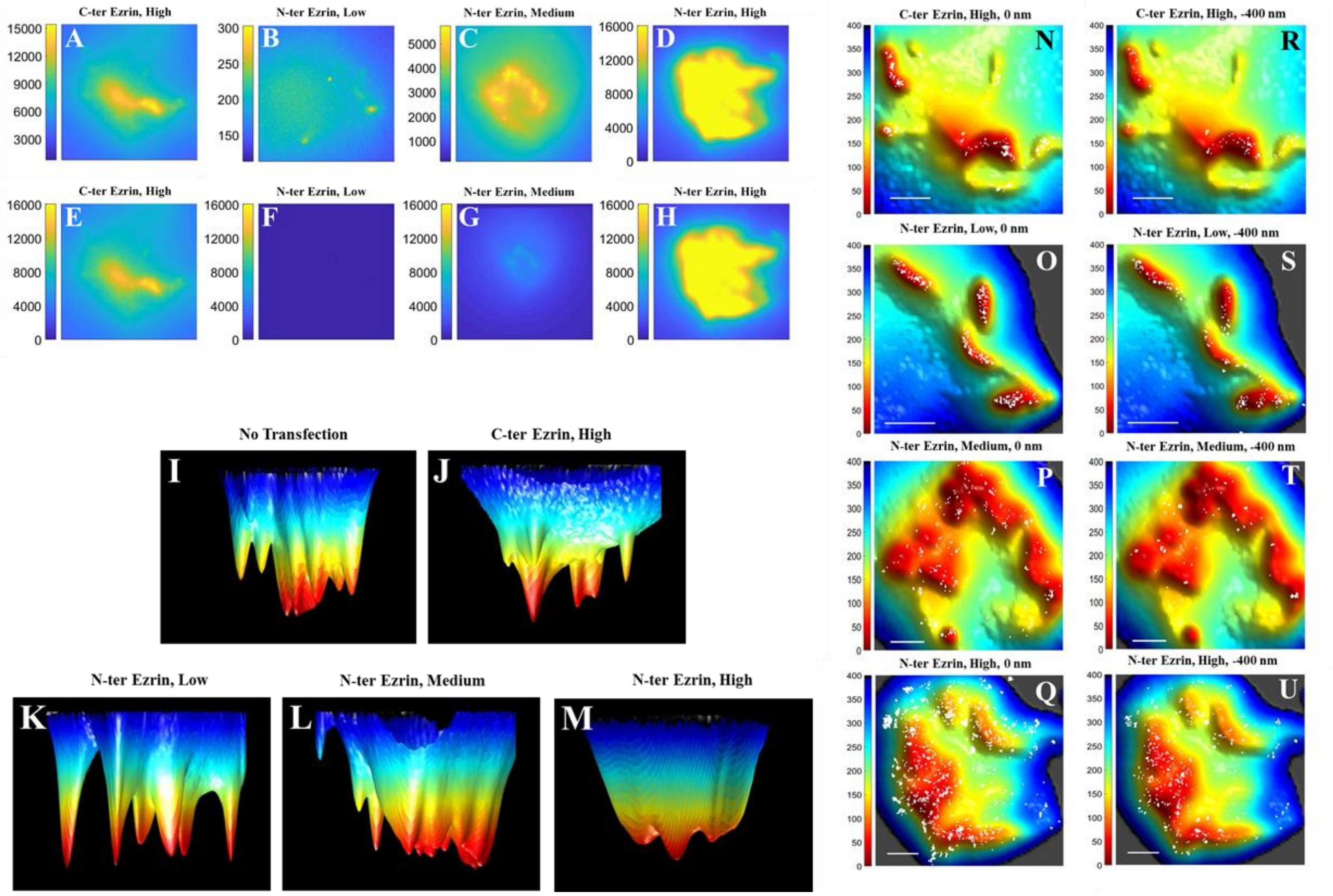


**Figure S5: Effect of dominant-negative ezrin transfection on the localization of TCRαβ with respect to 3D topography of Jurkat T cells (Related to Fig 7):**
Levels of Ezrin transfection measured through Alexa-405 fluorescence intensity. In A-D each image is normalized to the maximum intensity, in E-H the same images are normalized on a single scale.
Side views of membrane reconstruction images of Jurkat cells transfected with dominant-negative ezrin. Panels J-M correspond to the same panels in Figure S5A-D.
Super-resolution localization map (white dots) of Alexa-647-labelled TCRαβ at the (N-Q) 0 nm and (R-U) -400 nm plane are overlaid with the 3D surface reconstruction map of Jurkat cells transfected with various ezrin constructs. The color bars represent distance from the glass in nm. Scale bars: 1 µm.

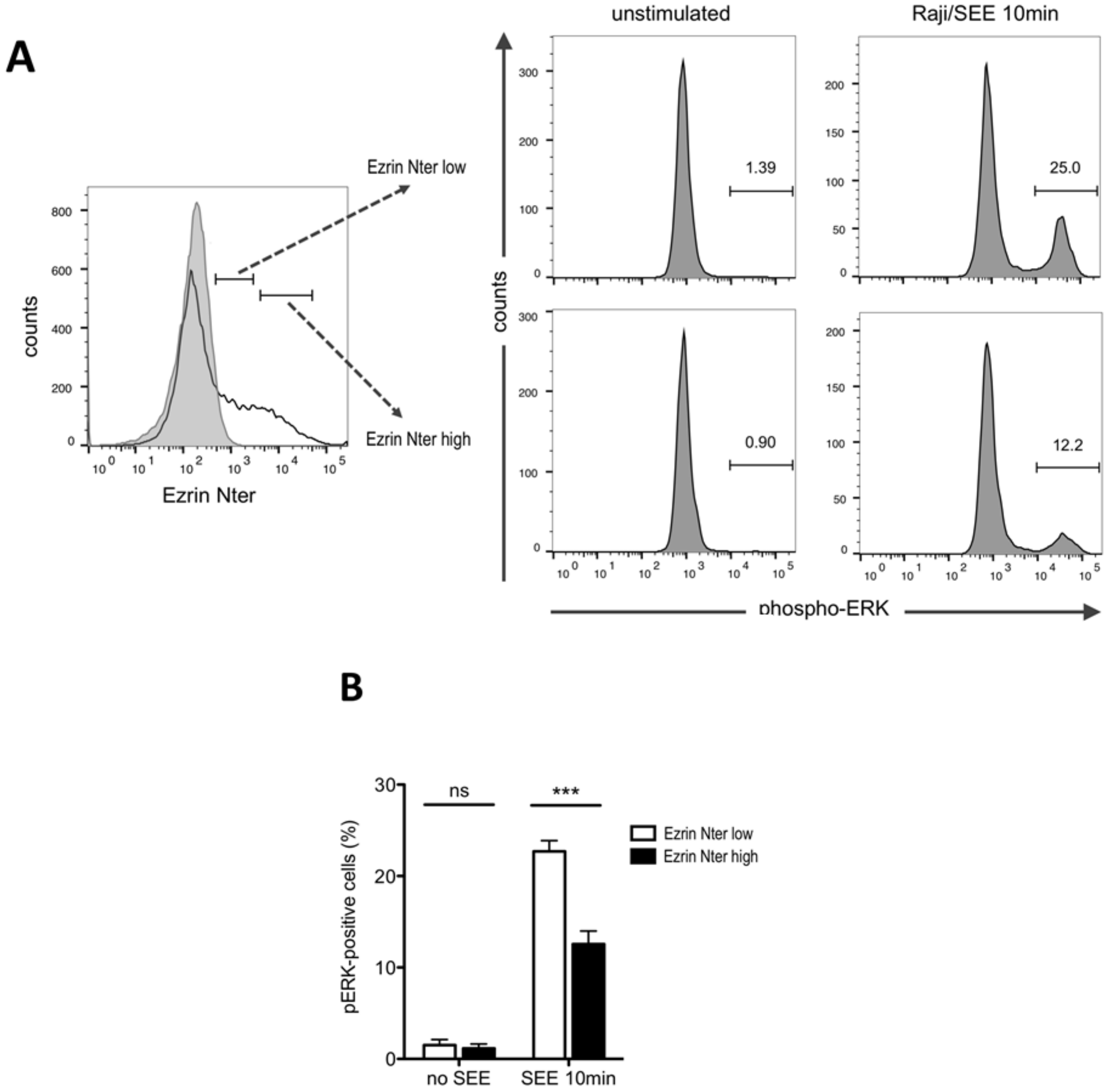


**Figure S6. ERK activation assay to address the functional consequences of altered distribution of TCRs in relation to microvilli (Related to Fig 7).** Jurkat cells transfected with N-ter ezrin expression plasmid or empty vector were incubated with Raji cells, alone or pre-loaded with SEE. After fixation, cells were permeabilized and stained with anti-VSV-G tag, to assess N-ter ezrin expression, and anti-phospho-ERK antibodies. A. Left: transfected Jurkat cells were gated based on the expression of low or high amounts of N-ter ezrin (left panel, black line). Gray-filled histogram shows labelling of control cells transfected with empty vector. Right: phospho-ERK staining of low (top row) or high (bottom row) N-ter ezrin-expressing cells, either unstimulated (Raji) or stimulated with SEE-loaded Raji (Raji+SEE). Numbers over the horizontal bars indicate the percentage of phospho-ERK-positive cells in each sample. The figure shows a representative experiment out of three performed. B. Quantification of phospho-ERK-positive cells in low and high N-ter ezrin expressing cells. Histogram show average ± SEM from three independent experiments. Statistical analysis was performed with a one-way ANOVA. ns: not significant difference. ***: $p<0.001$.

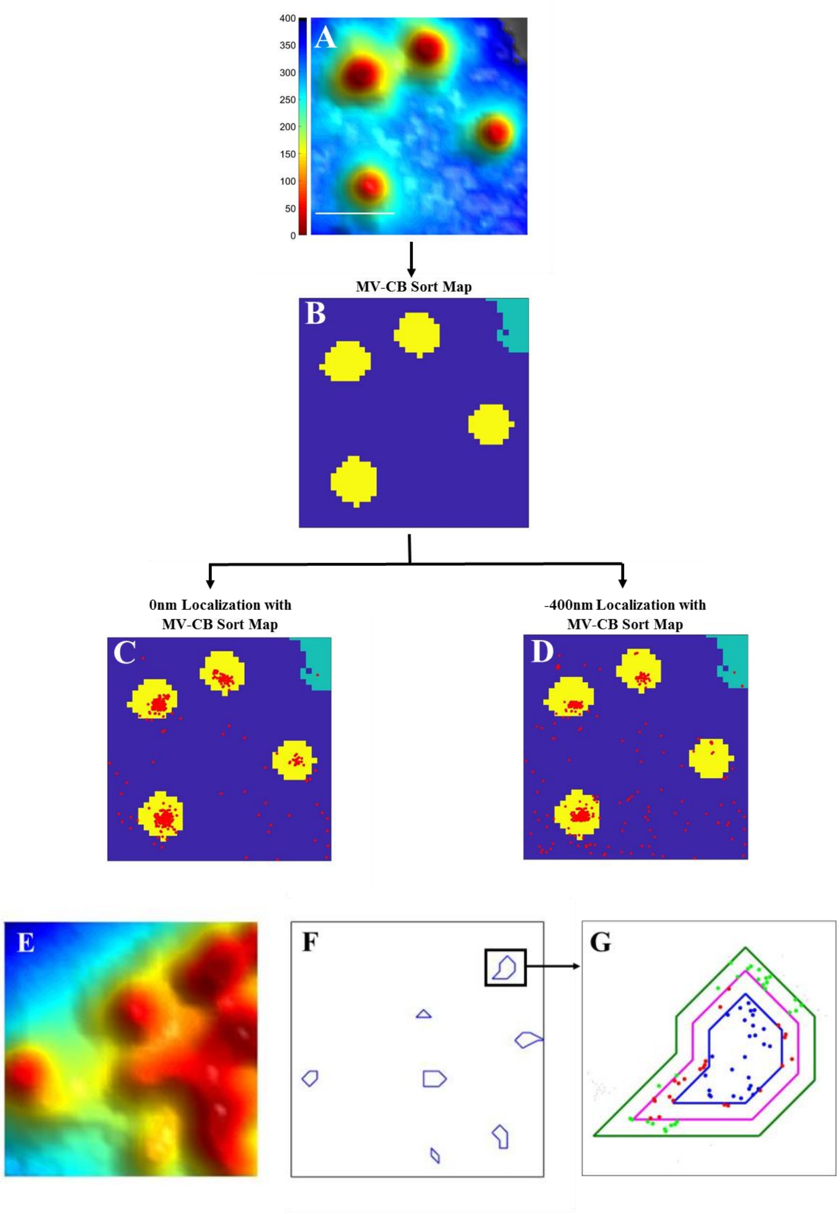


**Figure S7. Quantitative analysis of molecular distribution (Related to Fig. 4).**
Schematic representation of the method for sorting membrane proteins into microvilli (MV) and cell-body (CB) regions. (A) Representative 3D surface reconstruction map. Scale bar: 1 μm. (B) Microvilli (MV) and cell-body (CB) region segmentation map (MV: Yellow, CB: Blue, Background (BG): Green) (C, D) Localizations of a representative receptor molecules (red dots) obtained from the 0 nm plane (C) and the -400 nm plane (D) are overlaid with the segmentation map.
Schematic representation for determination of the cumulative fractional increase of the number of molecules of a specific protein on each cell as a function of the distance from the central region of each microvillus. (E) Membrane reconstruction image of a Jurkat cell. (F) Corresponding central microvilli regions marked with blue closed curves. (G) A representative plot of the concentric

shapes drawn around central microvilli regions to count the number of molecules as a function of the distance from the tips.

#### Supporting Tables

**Table S1: Fraction of molecules localized to microvilli**

| Receptor/adhesion molecule | No. of cells measured | Percentage of molecules on microvilli at the 0 nm Plane | Standard error of mean | Percentage of molecules on microvilli at the -400 nm Plane | Standard error of mean |
|---|---|---|---|---|---|
| TCRαβ | 9 | 89.0 | 2.4 | 86.1 | 2.2 |
| CD45 | 10 | 31.6 | 2.0 | 32.9 | 1.2 |
| CD4 | 10 | 94.4 | 1.1 | 83.8 | 2.6 |
| TCRζ | 11 | 96.0 | 1.2 | 96.2 | 1.2 |
| $Lck^{E}$ | 10 | 74.5 | 3.5 | 65.4 | 4.7 |
| $Lck^{J}$ | 10 | 76.3 | 4.1 | 59.6 | 4.8 |
| LAT | 11 | 67.8 | 2.9 | 54.6 | 2.3 |
| $CD2^{E}$ | 10 | 95.3 | 1.0 | 90.7 | 1.8 |
| $CD2^{J}$ | 10 | 91.4 | 1.8 | 85.2 | 2.4 |

The E and J superscripts following some protein names denote values obtained with effector cells and Jurkat cells, respectively. When no superscript is shown, the values were obtained with Jurkat cells.

**Table S2: Slopes of the δCount/δArea* of Figure 4B**

| Receptors/adhesion molecules | Slope | Standard error |
|---|---|---|
| TCRαβ | -0.155 | 0.002 |
| CD45 | 0.003 | 0.002 |
| TCR ζ | -0.101 | 0.004 |
| CD4 | -0.336 | 0.011 |
| LCK[E**] | -0.060 | 0.002 |
| LCK[J***] | -0.026 | 0.004 |
| LAT | -0.011 | 0.007 |
| CD2[E*****] | -0.072 | 0.009 |
| CD2[J******] | -0.090 | 0.006 |

The E and J superscripts following some protein names denote values obtained with effector cells and Jurkat cells, respectively. When no superscript is shown, the values were obtained with Jurkat cells.

* Calculated from 0-100 nm segment of the plot unless otherwise mentioned;

** Calculated from 30-100 nm segment of the plot

*** Calculated from 20-100 nm segment of the plot

***** Calculated from 40-100 nm segment of the plot

****** Calculated from 40-100 nm segment of the plot

### References


1 Jung, Y. *et al.* Three-dimensional localization of T-cell receptors in relation to microvilli using a combination of superresolution microscopies. *Proc Natl Acad Sci U S A* **113**, E5916-E5924, doi:10.1073/pnas.1605399113 (2016).

2 von Andrian, U. H., Hasslen, S. R., Nelson, R. D., Erlandsen, S. L. & Butcher, E. C. A central role for microvillous receptor presentation in leukocyte adhesion under flow. *Cell* **82**, 989-999 (1995).

3 Santos, A. M. *et al.* Capturing resting T cells: the perils of PLL. *Nat Immunol* **19**, 203-205, doi:10.1038/s41590-018-0048-8 (2018).

4 Fernandes, R. A. *et al.* A cell topography-based mechanism for ligand discrimination by the T cell receptor. *Proc Natl Acad Sci U S A* **116**, 14002-14010, doi:10.1073/pnas.1817255116 (2019).

5 Cai, E. *et al.* Visualizing dynamic microvillar search and stabilization during ligand detection by T cells. *Science* **356**, doi:10.1126/science.aal3118 (2017).

6 Majstoravich, S. *et al.* Lymphocyte microvilli are dynamic, actin-dependent structures that do not require Wiskott-Aldrich syndrome protein (WASp) for their morphology. *Blood* **104**, 1396-1403, doi:10.1182/blood-2004-02-0437 (2004).
